# A comprehensive analysis of gut and skin microbiota in canine atopic dermatitis in Shiba Inu dogs

**DOI:** 10.1101/2022.07.11.497949

**Authors:** Mirja Thomsen, Axel Künstner, Inken Wohlers, Michael Olbrich, Tim Lenfers, Takafumi Osumi, Yotaro Shimazaki, Koji Nishifuji, Saleh M Ibrahim, Adrian Watson, Hauke Busch, Misa Hirose

## Abstract

**Background:** Like its human counterpart, canine atopic dermatitis (cAD) is a chronic relapsing condition; thus, most cAD-affected dogs will require lifelong treatment to maintain an acceptable quality of life. A potential intervention is modulation of the composition of gut microbiota, and in fact, probiotic treatment has been proposed and tried in human atopic dermatitis (AD) patients. Since dogs are currently receiving intensive medical care, this will be the same option for dogs, while evidence of gut dysbiosis in cAD is still missing, although skin microbial profiling in cAD has been conducted in several studies. Therefore, we conducted a comprehensive analysis of both gut and skin microbiota in cAD in one specific cAD-predisposed breed, Shiba Inu. Additionally, we evaluated the impact of commonly used medical management on cAD (Janus kinase; JAK inhibitor, oclacitinib) on the gut and skin microbiota. Furthermore, we genotyped the Shiba Inu dogs according to the mitochondrial DNA haplogroup and assessed its association with the composition of the gut microbiota.

**Results:** *Staphylococcus* was the most predominant bacterial genus observed in the skin; *Escherichia/Shigella* and *Clostridium sensu stricto* were highly abundant in the gut of cAD-affected dogs. In the gut microbiota, *Fusobacteria* and *Megamonas* were highly abundant in healthy dogs but significantly reduced in cAD-affected dogs. The abundance of these bacterial taxa was positively correlated with the effect of the treatment and state of the disease. Oclacitinib treatment on cAD-affected dogs shifted the composition of microbiota towards that in healthy dogs, and the latter brought it much closer to healthy microbiota, particularly in the gut. Additionally, even within the same dog breed, the mtDNA haplogroup varied, and there was an association between the mtDNA haplogroup and gut microbial composition.

**Conclusions:** Dysbiosis of both the skin and the gut was observed in cAD in Shiba Inu dogs. Our findings provide a basis for the potential treatment of cAD by manipulating the gut microbiota as well as the skin microbiota.

## Background

Similar to humans, dogs suffer from spontaneously occurring diseases, such as cancers, and age-associated diseases [1–3]. Allergy is one such disease, and atopic dermatitis (AD) is the most common chronic inflammatory skin disorder both in humans and in dogs. In both species, AD is a complex and heterogeneous disease that is thought to be affected by multiple factors, such as host genetics and environmental factors [4–6]. In dogs, canine AD (cAD) exhibits similar clinical immunological characteristics to those of human AD [7]. cAD has a strong association with breed, although the breed predisposition varies among geographical locations [8], supporting a complex interaction of gene and environmental factors. Environmental factors, such as pH, temperature, hydration, hygiene practices, and food, play a pivotal role in maintaining the homeostasis of the host-microbiome [9], and changes in microbial composition, namely, dysbiosis, are known to drive the pathology of complex diseases [10]. In fact, dysbiosis of the skin microbiota has been the most well described in human AD patients, primarily defined as an overgrowth of *Staphylococcus spp.* on the human AD skin [11, 9]. Similarly, several studies have profiled skin microbiota in cAD-affected dogs, demonstrating skin dysbiosis in cAD [12–14]. Two of these studies have demonstrated a predominance of the genus *Staphylococcus* on the cAD dog skin compared to the healthy dog skin, confirming culture-based analysis of cAD skin that demonstrated the higher prevalence of this bacterial genus in cAD-affected dogs compared to healthy dogs [15].

Furthermore, a number of studies have demonstrated that gut dysbiosis is accompanied in human AD patients (reviewed in [16]). This is not surprising because both the gut and skin are abundantly colonized by distinct microbial communities, and these organs share a number of important anatomical and physiological characteristics [17]. In fact, although their relevance is underestimated and the mechanism is often not fully understood, an association between gut and skin manifestations has been reported in other pathological conditions in patients (reviewed in [17]). To date, the role of the gut microbiota has been described as an immune system regulator by its production of bacterial metabolites. The most well-known example is short-chain fatty acids (SCFAs), including butyrate, propionate and acetate, which have anti-inflammatory and immunomodulatory functions in other organs and the skin [18, 19]. As such, many studies suggest that probiotic supplementation may be a potentially effective treatment option for human AD; however, further studies are needed [20]. This is considered to be a similar situation for the clinical management of cAD. However, to the best of our knowledge, no study dissecting the gut microbiota in cAD-affected dogs has been reported thus far.

This is the first study that describes a comprehensive microbiome profile of both skin and faeces obtained from cAD and healthy Shiba Inu dogs. This dog breed is highly susceptible to cAD in Japan, as shown by a high number of cases in published studies of cAD in this region [21–23]. More specifically, we analysed the microbiota composition of twelve skin sites, which were evaluated for clinical score (the fourth version of the Canine Atopic Dermatitis Extent and Severity Index; CADESI-04), as well as intestinal microbiota in faecal samples of healthy and cAD Shiba Inu dogs. In addition, we were able to evaluate the impact of Janus kinase (JAK) inhibitor oclasitinib (Apoquel^®^; Zoetis, Parsippany, NJ, USA) treatment, which is commonly used for cAD treatment, on the gut and skin microbiota of cAD-affected dogs. We focused on one specific dog breed to minimize the potential influence of the host nuclear genome on the skin and intestinal microbial community between samples. Meanwhile, the impact of the mitochondrial DNA (mtDNA) has not been evaluated in this dog breed, particularly in the context of microbiota as well as pathological relevance. Since our group and others have demonstrated that variations in mtDNA are associated with the composition of microbiota in humans and rodents [24, 25], we sequenced the whole mitochondrial genome of Shiba Inu dogs to evaluate the potential association between mtDNA variants and microbiota composition in cAD-affected dogs.

## Methods

### Sample subjects and sampling for microbiome analysis and mitochondrial genome sequencing

In total, 40 dogs were recruited, of which 20 were healthy dogs (Healthy). Shiba Inu dogs diagnosed with cAD at Animal Medical Centre, Faculty of Agriculture, Tokyo University of Agriculture and Technology and private veterinary practices in Japan were recruited for this study. All dogs with cAD fulfilled more than five of Favrot’s diagnostic criteria, and other potential causes for pruritis, such as ectoparasite infestations and bacterial or fungal infections were excluded, according to a detailed guideline [26]. CADESI-04 scores for the same case were monitored by the same veterinary clinicians (TO, and YS). The Pruritis Visual Analog Scale (PVAS) was graded by participating cAD-affected dog owners to assess the severity of pruritic manifestation [27]. The clinical scores (CADESI-04 and PVAS) of cAD-affected dogs are presented in **Fig. S1**.

Of the 20 cAD dogs, 10 dogs were newly diagnosed with cAD without any previous medical treatments (cADpre), and 10 were already diagnosed with cAD before the study cohort recruitment and had been on oclacitinib treatment (cADtreat). The age and sex of the recruited dogs are summarized in **Table S1**. The ten dogs newly diagnosed with cAD (cADpre) were consequently under oclacitinib treatment for two weeks after the diagnosis. After the two-week treatment, the dogs were clinically re-evaluated and skin swab and faecal samples were obtained (cADpost). cADtreat dogs had been on oclacitinib treatment for an average of 13.24 months (standard deviation, s.d. ±8.458). Additionally, 20 healthy Shiba Inu dogs were recruited upon their visits for annual vaccination or general health checks. The dog groups tested in this study is depicted in **Fig. S1a**. Clinical history, housing environment, and dietary information were collected and recorded (**Supplementary Material S1**).

Skin swab samples were obtained from the perilesional area of 12 body sites, which are included in the CADESI-04 evaluation. All dogs did not receive topical disinfectants or shampoos at least three days before the sampling. Samples were collected from the left side of the body sites unless skin lesions were present only on the right side; since there were no cases of unilateral right-sided lesions, all samples were collected from the left body side. Sterile culture swab applicators were soaked in SCF-1 buffer (50 mM Tris, 1 mM EDTA, 0.5% Tween 20; Teknova, CA, USA) and rubbed on the perilesional skin 40 times, while the swab was rotated every 10 strokes. Swabs were stored in tubes filled with 400 µl of SCF-1 buffer at −80 °C until bacterial DNA isolation was performed. Faecal samples were also collected from the dogs by inserting the swab into the rectum and stored in tubes filled with 400 µl SCF-1 buffer at −80 °C. A total of 600 skin samples, 50 faecal samples, and 40 buccal samples were collected from the study cohort.

### Bacterial DNA isolation, library preparation and sequencing of the bacterial 16S rRNA gene

*Skin microbial DNA isolation, library preparation, and sequencing*: Microbial DNA was extracted from the skin swab samples using a QIAmp DNA Microbiome Kit (Qiagen, Hildfeld, Germany) according to the manufacturer’s protocol. Skin microbial DNA samples were further processed for library preparation for the hypervariable V1-V2 region of the 16S rRNA gene, as previously described [25,28,29]. The final library was sequenced on the Illumina MiSeq platform using v2 chemistry, generating 2 × 250bp reads.

*Faecal microbial DNA isolation, library preparation, and sequencing:* Microbial DNA was prepared from the faecal samples using a Qiagen Power Soil Kit (Qiagen) according to the manufacturer’s protocol. Faecal microbial DNA samples were further processed for library preparation for the 16S rRNA gene. The 16S rRNA gene was amplified using uniquely barcoded primers flanking the V3 and V4 hypervariable regions (341F - 806R) with fused MiSeq adapters and heterogeneity spacers in a 25 µl volume, consisting of one µl template DNA, four µl of each forward and reverse primer, 0.25 µl Phusion Hot Start II DNA polymerase (0.5 units), 0.5 µl dNTPs (200 µM each) and five µl of HF buffer. The PCR conditions were as follows: initial denaturation for 30 sec at 98 °C; 32 cycles of 9 sec at 98 °C, 30 sec at 55 °C, and 45 sec at 72 °C; and a final extension for 10 min at 72 °C. The concentration of the PCR products was estimated on 1.5% agarose gels using the image software Quantum Capt v16.04 (Vilber Lourmat Deutschland GmbH, Eberhardzell, Germany) with a known concentration of DNA marker (100 bp DNA ladder; Thermo Fisher Scientific GmbH, Dreieich, Germany) as the internal standard for band intensity measurement. The PCR products were pooled into approximately equimolar subpools, as indicated by band intensity, followed by clean-up of the subpools using a GeneJet column purification kit (Thermo Fisher). The concentrations of subpools were quantified using a Qubit dsDNA BR Assay Kit on a Qubit fluorometer (Thermo Fisher Scientific GmbH). The subpools were combined in one equimolar final pool for each library. The final pools were cleaned up with magnetic beads (MagSi-NGS^PREP^ Plus; Steinbrenner Laborsysteme GmbH, Wiesenbach, Germany), the concentration was measured by qPCR using a NEBNext Library Quant Kit for Illumina (New England BioLabs GmbH, Hessen, Germany), and the size was determined by Bioanalyzer 2100 using Agilent High Sensitivity DNA Kit (Agilent Technologies GmbH, Waldbronn, Germany). The final library was sequenced on the Illumina MiSeq platform using v3 chemistry, generating 2 × 300bp paired-end reads (Illumina).

### Buccal DNA isolation, library preparation and sequencing of the whole mitochondrial genome

*Genomic DNA isolation from buccal swab samples*: Genomic DNA was isolated from the buccal swab samples using a DNeasy Blood/Tissue Kit (Qiagen) according to the standard protocol. Genomic DNA (gDNA) samples were processed for library preparation, as previously described in the Human mtDNA Genome protocol for Illumina Sequencing Platform [30], with small modifications. In brief, two primer sets [CLF_mtDNA_1_F (TTCTTCGGCGCATTCCACAA) and CLF_mtDNA_1_R (GGCATGCCTGTTAATGCGAG); CLF_mtDNA_2_F (TCACTTTATGCTTAGGGGCCA) and CLF_mtDNA_2_R (CAACATTTTCGGGGTATGGGC)] were used to enrich the canine mtDNA by long-range PCR. These PCR products cover the whole canine mtDNA sequencing (16,727 bp). The PCR reagents were combined in a total reaction volume of 20 µl, consisting of 10 ng template gDNA, 10 µM of each forward and reverse primer, 10 mM dNTPs, Phusion High-Fidelity DNA polymerase (Thermo Fisher, Germany) and 5× Phusion HF buffer. The PCR conditions for the amplification of the CLF_mtDNA_1 fragment were 98 °C for 30 sec; 29 cycles of 98 °C for 5 sec, 67.7 °C for 10 sec, and 72 °C for 4 min 45 sec; 72 °C for 5 min; hold at 4 °C, and those for the amplification of the CLF_mtDNA_2 fragment were 98 °C for 30 sec; 29 cycles of 98 °C for 5 sec, 65.8 °C for 10 sec, and 72° C for 4 min 15 sec; 72 °C for 5 min; and hold at 4 °C. Library preparation was performed using a Nextera XT DNA Library Preparation Kit (Illumina Inc., CA, USA) according to the manufacturer’s instructions. The 10-pM final library was sequenced on the Illumina MiSeq sequencing platform (2 × 150bp paired-end reads) (Illumina Inc.).

### Microbiota data processing

Raw sequencing reads in fastq format were merged and filtered for low quality (VSEARCH [31] v2.12.0; maxdiffs = 2, maxee = 0.5), and chimeric sequences were removed from the data (VSEARCH with RDP Gold v9 as reference database). Afterwards, de-replication was performed (VSEARCH) and sequences were clustered into operational taxonomic units (OTUs) using UPARSE [32] as implemented in USEARCH (v11.0.667). Taxonomic assignment was performed applying the SINTAX [33] algorithm (vsearch) with RDP v18 [34] as the classification database. Chimaera-free merged reads were mapped to OTUs using VSEARCH (command: *usearch_global*) with a 97% identity threshold to create the OTU table. The phylogenetic tree was reconstructed by aligning OTU sequences using MOTHUR [35] (v.1.41.3) and SILVA [36] v123 as a reference database. Afterwards, tree reconstruction was performed by using FASTTREE [37] (v2.1) with a generalized time-reversible model with gamma correction. OTUs belonging to the genus *Staphylococcus* were further specified at the species level using NCBI Web BLAST (Nucleotide BLAST; https://blast.ncbi.nlm.nih.gov/Blast.cgi) against the 16S rRNA database with default parameters.

*Microbiome data filtering*: The data were imported into R (v4.2.0) using phyloseq [38] (v1.40.0), and covariates were matched. OTUs not matching *Bacteria* or unknown phylum were excluded. Additionally, OTUs matching the phyla *Abditibacteriota*, *Armatimonadetes*, *Balneolaeota*, *Chloroflexi*, *Cyanobacteria/Chloroplast*, *Elusimicrobia*, *Gemmatimonadetes*, *Latescibacteria*, *Nitrospirae*, *Planctomycetes*, or *Rhodothermaeota* were excluded as potential contaminants. Next, decontam [39] (v1.16.0), a contamination detection tool, was applied to remove contaminants based on frequency (threshold 0.1; 114 OTUs excluded after manual inspection of potential contaminants) and prevalence (threshold 0.2; 72 OTUs excluded). Finally, only samples reaching at least 2,300 contigs were included in the downstream analysis (7 samples excluded).

*Alpha and beta diversity*: In faecal samples, alpha diversity was estimated using DivNet [40] (v0.4.0; EMiter = 6, EMburn = 3, MCiter = 250, MCburn = 100, network = diagonal). To test for significant differences in DivNet-estimated Shannon diversity, we tested for heterogeneity of total diversity (observed plus unobserved) across multiple sites using the *betta* [41] function (breakaway v4.7.9). Groupwise estimates were compared using the *testDiversity* function (DivNet). For skin samples, alpha diversity (Shannon index) was estimated using the *estimate_richness* function of phyloseq, and significance was assessed using non-parametric tests (Wilcoxon test for pairwise testing and Kruskal–Wallis test for more than 2 group comparisons with pairwise Wilcoxon tests as *post-hoc* test).

Beta diversity was estimated using centred log-ratio transformed (*clr*) data and distances were calculated using Euclidean distance (Aitchison distance [42]). Permutational multivariate analysis of variance using distance matrices (PERMANOVA) was used to analyse differences in beta diversity (*adonis2* function, as implemented in the vegan package v2.6-2, and *pairwise.perm.manova* function, as implemented in the RVAideMemoire package v0.9.79, each with 99,999 permutations). Inter-object dissimilarities were visualized using constrained analysis of principal coordinates (CAP) (vegan package v2.6-2).

*Differentially abundant taxa*: Differentially abundant (DA) taxa were identified on prevalence filtered data (taxa with a prevalence <20% were excluded in each comparison). A count regression for correlated observations with a beta-binomial distribution model as implemented in the corncob [43] package (v0.2.0) was applied, and taxa with a *q*-value below 0.1 were considered significant. To correct for sex differences in the analysis, sex was incorporated into the NULL model in the intra-location comparisons (*H_0_ ∼ sex*, *H_1_ ∼ group + sex*). For the haplogroup analysis, disease group was incorporated into the NULL model to correct for these effects (*H_0_ ∼ group*, *H_1_ ∼ group + haplogroup*). To verify the findings, ANCOM-BC [44] (v1.6.0) was used (lib_cut = 0, struc_zero = TRUE, neg_lb = FALSE, tol = 1e-5, max_iter = 10000), using the same models as in the case of corncob. Again, taxa with a *q*-value below 0.1 were considered significantly different. Differential abundance testing for *Staphylococcus spp.* was performed using ANCOM-BC.

*Statistical analysis*: If not stated differently, analyses were performed using R (v4.2.0), and *p* values were corrected for multiple testing using Benjamini–Hochberg correction (denoted using *q* or *fdr*). For data handling, phyloseq and tidyverse (v1.3.1) were used; cowplot (v1.1.1), ggpubr (0.4.0), sjplot (v2.8.10) and patchwork (v1.1.1) were applied to create the figures. Balances with CADESI-4 or PVAS as the target variable were calculated using a forward-selection method as implemented in selbal [45] (v0.1.0) on phyla/genera occurring in at least one-third of the samples per location with 5-fold cross-validation (10 iterations each).

For correlation analysis (Spearman rank correlation), *clr*-transformed abundance values were used. At the genus level, only genera found in at least 1/15th of the samples were included to calculate correlations.

The workflow used for the microbiota analysis is available at https://github.com/buschlab/2022_canine_atopic_dermatitis_paper.

### Mitochondrial genome analysis

Raw sequencing data were mapped with bwa (version 0.7.17) [46] against dog reference sequence U96639.2 (https://www.ncbi.nlm.nih.gov/nuccore/U96639), and variant calling was performed with bcftools (version 1.15) [47] using parameter --ploidy 1, i.e., assuming a ploidy of 1. The PHRED-scaled genotype likelihood was visualized using a heatmap. This value ranges between 0 and 255, with 0 being the strongest support of a homomorphic non-reference (alternative) allele. Values larger than 0 were typically seen at positions at which more than one allele is observed, indicative of variant calling artefacts or heteroplasmic variants. Variants with no genotype calls, i.e., homomorphic reference allele, are displayed in white. Mitochondrial (MT) sequences were generated by removing indels and variants with quality less than 200 and then applying the remaining variants to the reference sequence using the bcftools consensus. The resulting MT sequences have individually been uploaded to the Canis mtDNA HV1 database (http://chd.vnbiology.com) [48] for haplogroup assignment. The MT analyses workflow used is available at https://github.com/TLenfers/multispecies_mitochondrial_variant_analysis. Furthermore, MT sequences used for haplogroup assignment as well as the code for generating Figure S11, including underlying variant data, are available at https://github.com/iwohlers/2022_dog_mt.

## Results

### Differential microbiota composition in healthy and cAD-affected dogs with and without medical treatment

Five hundred fifty-four skin swab samples were successfully sequenced for the hypervariable V1-V2 region of the 16S rRNA gene. On average, 44,387 (s.d. ±26,307) read pairs were obtained in each sample. Merging forward and reverse reads into contigs, quality filtering, and chimaera removal resulted in an average of 29,343 (±19,755) contigs per sample.

Taxonomic classification of 12 skin sites from healthy dogs, untreated cAD-affected dogs (cADpre), the same cADpre dogs that received 2 weeks of oclacitinib treatment (cADpost), and cAD-affected dogs that had already been on the treatment (cADtreat) dogs at the phylum and genus levels are shown in **Fig. 1a** and **Fig. 1b** (bacterial taxa identified in these skin samples are shown in **Supplementary Material S2** as well). While there is an obvious heterogeneity in the relative abundance of the skin microbiota across the 12 skin sites, *Staphylococcus* was the predominant bacterial taxon in all 12 skin sites of cAD-affected dogs.

**Fig. 1.**
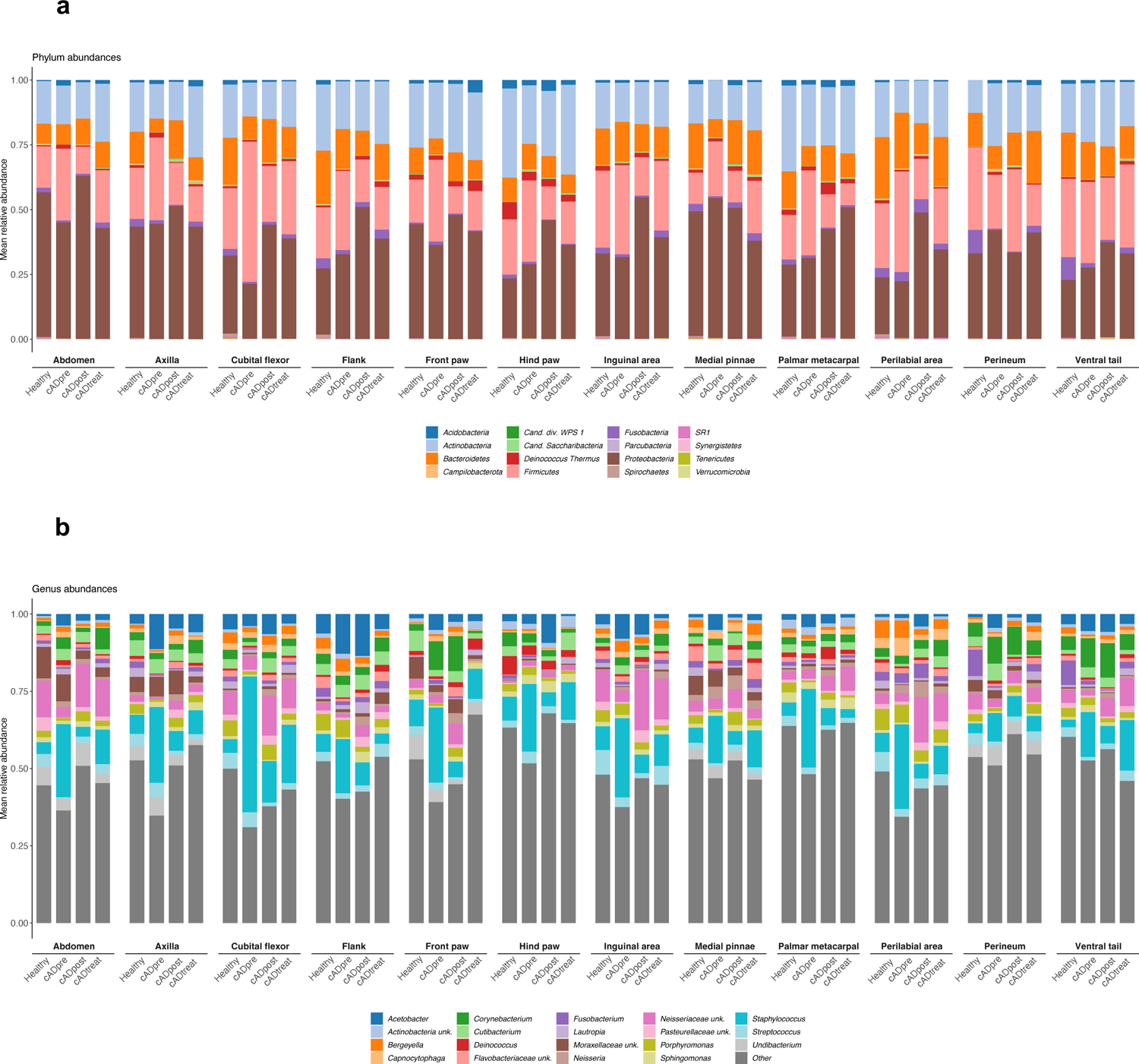
Taxonomic classification of 12 skin sites from healthy dogs, dogs with untreated cAD (cADpre), cAD-affected dogs that received 2 weeks of oclacitinib treatment (cADpost) and cAD-affected dogs that had already been on the treatment (cADtreat) at the phylum level (**a**) and the genus level (**b**).

### Reduction of bacterial diversity in cAD skin

We selected the skin sites that were most relevant to the clinical score for further analysis. To select these skin sites, balances of bacterial taxa were calculated using a greedy stepwise algorithm (selbal) with 5-fold cross-validation and 10 iterations. This method estimates the optimal number of discriminating taxonomic groups and the ratio of these taxonomic groups (named “denominator” and “numerator”) and then defines the balance between the microbial characteristics that best describe the differences in the compared phenotype — in this case, CADESI-04. The skin sites whose bacterial taxa at the genus level significantly correlated with their CADESI-04 scores (*R^2^* > 0.4) were selected for further analysis (**Table S2**). To this end, skin microbiome data from four skin sites, namely, the abdomen, front paw, perilabial area, and ventral tail, were selected for more detailed analysis.

Skin microbiome data of the four selected skin sites were analysed for alpha diversity (**Fig. S2**), and the difference in skin bacterial community composition was evaluated by beta diversity (**Fig. S3**). Alpha diversity (estimated using Shannon diversity) showed a general trend of reduction of bacterial richness in untreated cAD-affected dogs (cADpre) compared with healthy dogs, and the level was recovered after the medical intervention. A significant difference was observed in the perilabial area (*p* = 0.023), with a significant reduction in cADpre dogs (*p* = 0.011) and cADpost dogs (*p* = 0.027) compared with healthy dogs. Beta diversity was significantly different between groups in abdomen, front paw, and ventral tail (**Fig. S3**).

### Differentially abundant bacterial taxa in the skin between healthy and cAD-affected dogs with and without oclacitinib treatment

Next, for the four selected skin sites, differentially abundant bacterial taxa at the phylum level were analysed between two groups: cADpre *vs.* healthy, cADtreatment *vs.* healthy, and cADpost *vs.* cADpre (**Fig. S4** and **Table S4**). The abundance of *Firmicutes* in the front paw and perilabial area was significantly higher in cAD-affected dogs without treatment than in healthy dogs (cADpre *vs.* healthy). At these skin sites, cAD-affected dogs showed a significant reduction in *Firmicutes* abundance after 2 weeks of oclasitinib treatment (cADpost *vs.* cADpre). At all four skin sites, the abundance of *Proteobacteria* was significantly increased in cAD-affected dogs after they received 2 weeks of oclacitinib treatment (cADpost *vs.* cADpre). In contrast, the abundance of *Deinococcus-Thermus* significantly decreased after treatment in the abdomen, front paw, and ventral tail. Similarly, differentially abundant bacterial taxa at the genus level were analysed between the same two groups (**Fig. S5** and **Table S4**). The abundance of *Staphylococcus* was significantly higher in the perilabial area and ventral tail of cAD-affected dogs than in the same skin sites of healthy dogs. When compared with paired samples before and after the 2-week oclacitinib treatment (cADpost *vs.* cADpre), the abundance of *Staphylococcus* in the perilabial area was largely decreased after the treatment (*q* < 1.00×10^-50^, effect size = −2.4017), while the change in abdomen, front paw, and ventral tail was marginal (*q* = 3.50×10^-41^, effect size = 0.8125; *q* = 1.64×10^-6^, effect size = 0.1715; *q* = 2.25×10^-8^, effect size = −0.3779, respectively). Interestingly, the abundance of *Staphylococcus* on the abdomen of cAD-affected dogs that already received oclacitinib treatment (cADtreat), was less than that in the site-matched skin of healthy dogs. Meanwhile, there was still a higher bacterial load on the ventral tail of cADtreat dogs than in the same skin site on healthy dogs (*q* = 0.0554, effect size = 1.5864).

### Correlation analysis between bacterial taxa and clinical cAD parameters

Next, we conducted a correlation analysis between clinical parameters (CADESI-04 and PVAS) and microorganisms on the skin in each group to evaluate whether identified differences in the abundance of bacteria were associated with the clinical parameters (**Fig. 2** and **Fig. S6**, respectively). To identify specific bacterial phyla and/or genera that correlate with respective clinical phenotypes and are commonly shared in all cAD-affected dogs, we looked for similar trends of correlation in all three dog groups in all four skin sites. More specifically, bacterial taxa showing similar colours across the three groups in each heatmap were evaluated. There were no bacterial phyla and/or genera that had a similar impact (i.e., negative or positive relations) in CADESI-04 or PVAS. Only two genera, *Corynebacterium* and *Staphylococcus,* demonstrated a trend of positive correlation with CADESI-04 values in all groups in all four skin sites (**Fig. 2b**).

**Fig. 2.**
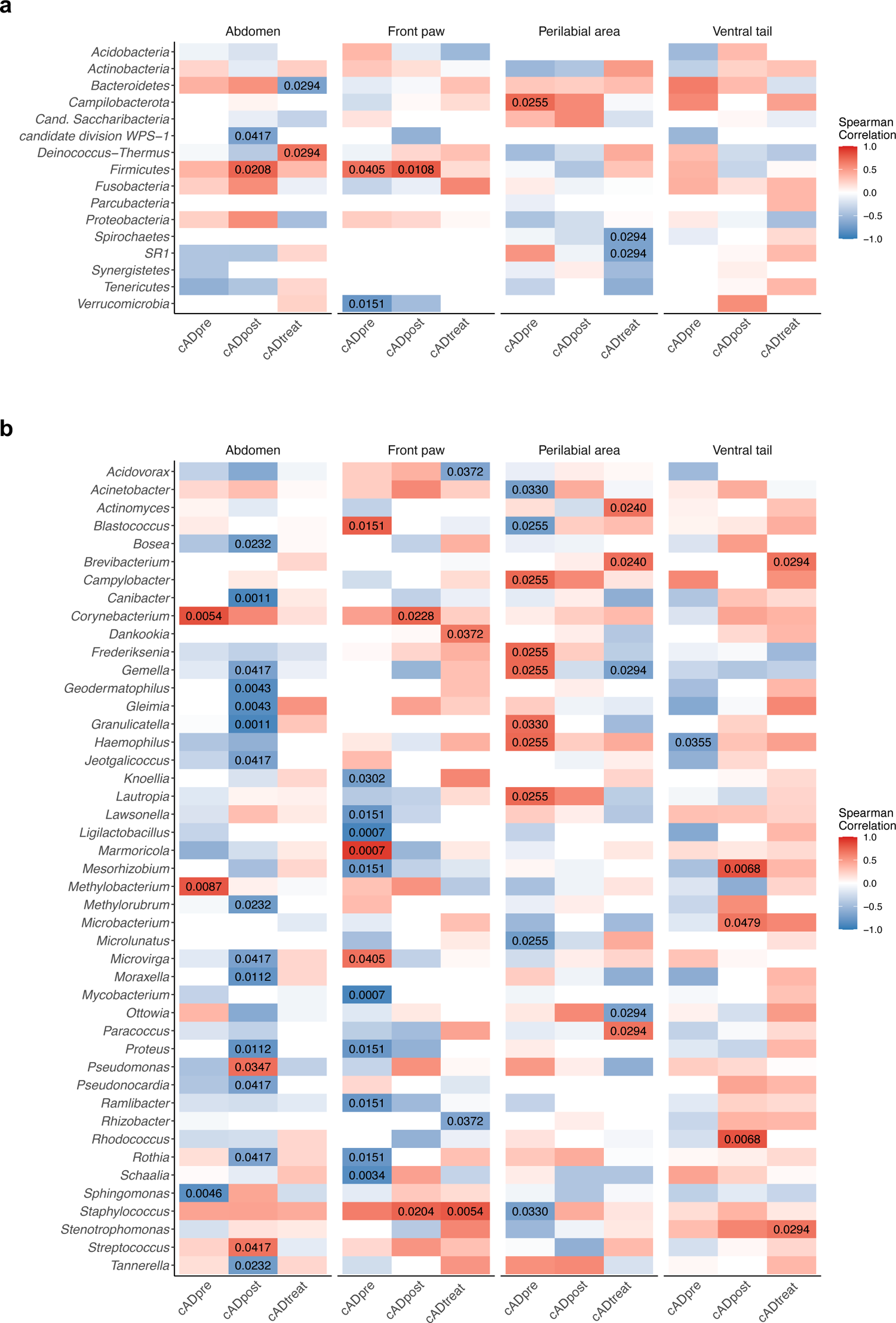
Spearman’s correlation of clinical score (CADESI-04) and bacterial taxa at the phylum level (**a**) and at the genus level (**b**) at the four skin sites: abdomen, front paw, perilabial area, and ventral tail. P values showing significant correlation (*p* < 0.05) are shown.

### Association of *Staphylococcus* abundance with cAD

Having our correlation analysis as well as previously reported clinical evidence, we further investigated whether the abundance of *Staphylococcus* was associated with cAD in all 12 skin sites (**Fig. S7** and **Table S4**). Of these skin sites, five (namely, the abdomen, axilla, cubital flexor, palmar metacarpal, and perilabial area) showed a significant association of *Staphylococcus* abundance between cADpre dogs and healthy dogs (*p* = 0.0392, *p* = 0.0436, *p* = 0.0026, *p* = 0.0105, and *p* = 0.0233, respectively; **Fig. S7a-e**). Even though the *p* values were not statistically significant (due to the large variation between individuals), the abundance of *Staphylococcus* in the other three skin sites (hind paw, flank, and front paw) showed a trend towards higher abundances in cADpre dogs than in healthy dogs (*p* = 0.2331, *p* = 0.1221, and *p* = 0.2889, respectively; **Fig. S7f-h**). These skin sites demonstrated a positive correlation with the corresponding CADESI-04 value (**Fig. S7i-p**).

Of the identified OTUs, fourteen different OTUs were mapped to the genus *Staphylococcus*. These OTUs were further classified at species level by an NCBI Nucleotide BLAST search. The ten most abundant species are presented in **Fig. S8** and **Table S4**. In each of the four selected skin sites (abdomen, front paw, perilabial area, and ventral tail), the abundance of *S. pseudintermedius* was significantly higher in cAD-affected dogs not receiving medical treatment (cADpre) than in healthy dogs (*q* < 0.1; **Fig. 3**).

**Fig. 3.**
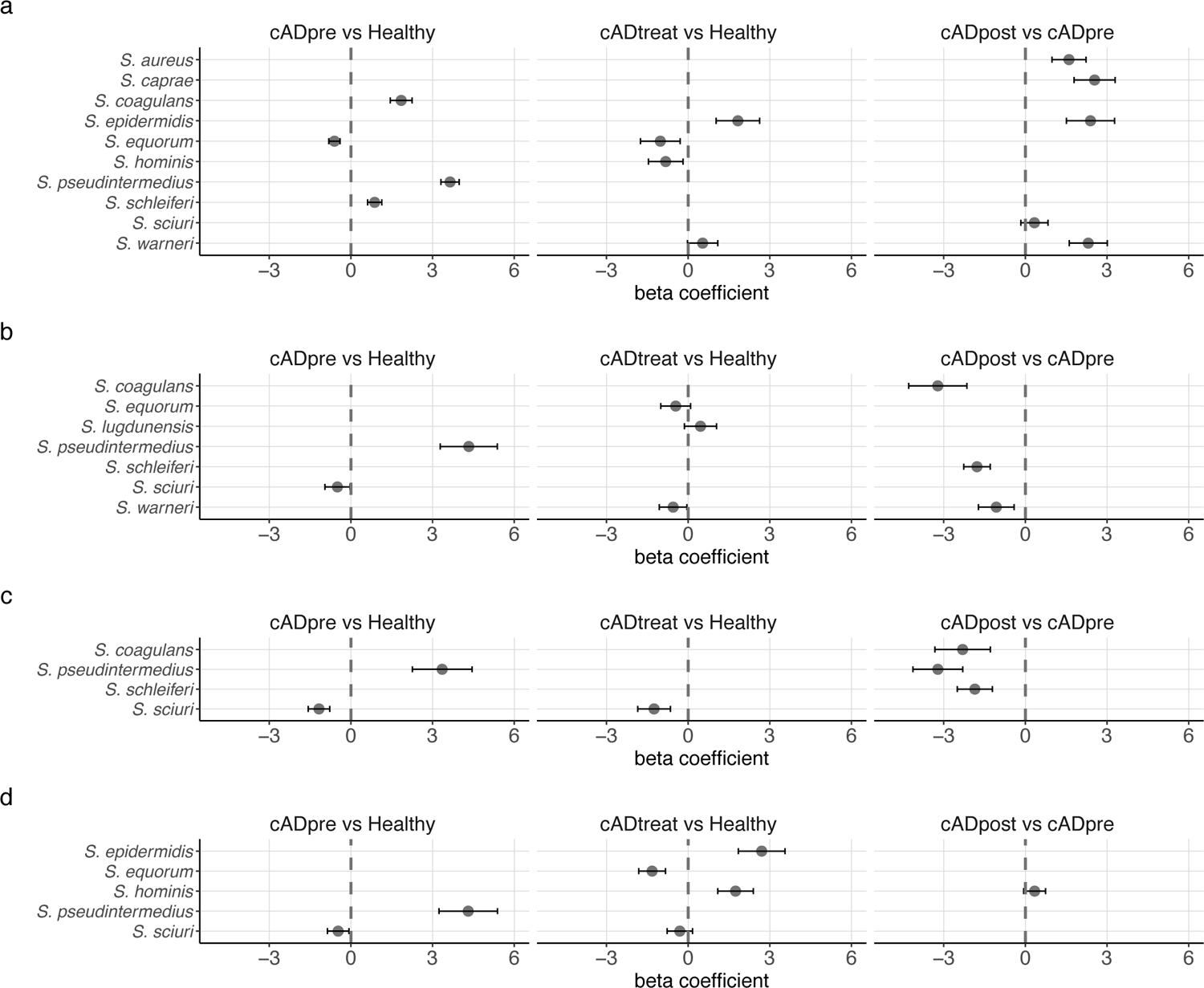
The abundances of the 10 predominant *Staphylococcus* spp. were compared between the two groups: cADpre *vs.* healthy, cADtreat *vs.* healthy and cADpost *vs.* cADpre at the four skin skites: abdomen (**a**), front paw (**b**), perilabial area (**c**), and ventral tail (**e**). The beta coefficient represents the coefficients obtained from the ANCOM-BC log-linear (natural log) model, and the standard error of the coefficients are shown as whiskers.

### Distinct gut microbial composition between healthy and cAD-affected dogs with and without oclacitinib treatment

A total of 46 faecal samples were analysed. Approximately 41,000 (± 25,895) read pairs per sample were obtained, and merging forward and reverse reads into contigs yielded approximately 41,000 (± 21,305) contigs per sample. After filtering and chimaera removal, the average number of contigs per sample was 36,367 (± 17,293).

The bacterial taxonomic composition in the faecal samples from all dogs was plotted (**Fig. 4**). The mean relative abundances of bacterial taxa in the four groups were investigated. A total of ten phyla were identified in the faecal samples (**Fig. 4a**). *Firmicutes* was the most abundant, followed by *Bacteroidetes* and *Proteobacteria*. We observed a major decrease in the proportion of *Fusobacteria* accompanied by an increase in *Proteobacteria* in all cAD groups (cADpre, cADpost, and cADtreat) (**Table S5**).

**Fig. 4.**
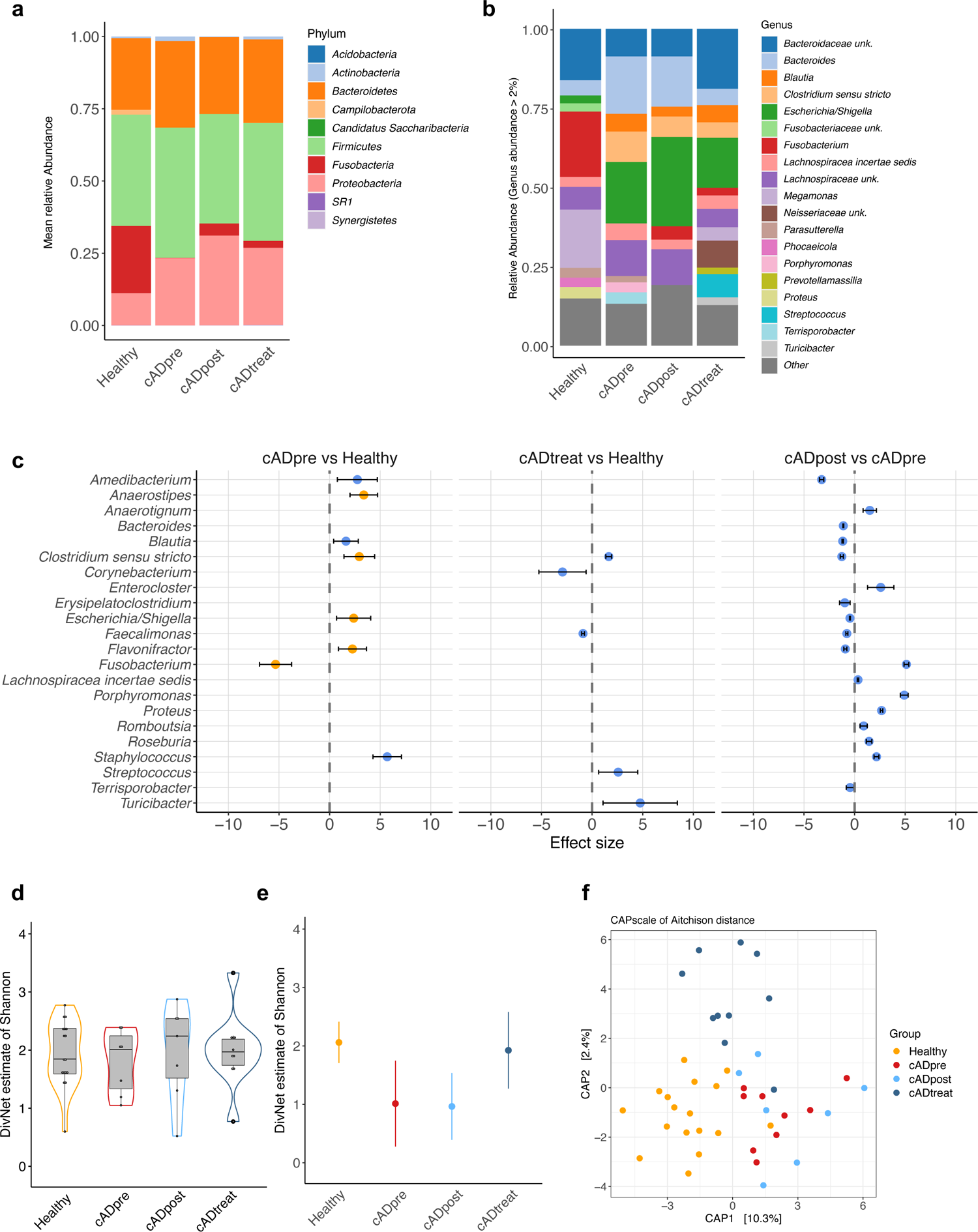
Faecal microbiota analysis. Mean relative taxonomic abundances at the phylum level (**a**) and at the genus level (**b**) in each dog group are presented. The differentially abundant bacterial genera (*q* < 0.1) are presented (**c**), where blue dots refer to genera identified by corncob and orange dots to genera identified by corncob and ANCOM-BC. The comparison of the abundance was conducted between the two groups indicated in each panel: left, cAD-affected dogs that did not receive medical treatment (cADpre) *vs.* healthy dogs (Healthy); middle, cAD-affected dogs on medical treatment (cADtreat) *vs.* healthy dogs; right, cADpre dogs that received 2 weeks of oclacitinib treatment (cADpost) *vs*. cADpre. Alpha diversity was analysed using the DivNet estimate of Shannon, as samplewise estimates (**d**) and communitywise estimate (**e**). Beta diversity was evaluated using constrained analysis of principal coordinates of the Aichison distance, demonstrating clear differences between the four groups (adonis: *p* < *0.001*).

A total of 131 different genera were identified in the faecal samples. The mean relative abundance of the 19 most abundant genera is shown in **Fig. 4b**. In healthy dogs, the predominant genera were *Fusobacterium* (20.6%), *Megamonas* (18.4%), and an unclassified genus belonging to the *Bacteroidaceae* family (16.2%). In contrast, the predominant genera found in untreated cAD-affected dogs (cADpre) were *Escherichia*/*Shigella* (19.4%), *Bacteroides* (18.1%), and *Clostridium sensu stricto* (9.7%), with *Fusobacterium* (0.06%) and *Megamonas* (0.0003%) being detected at very low abundances in this group (**Table S6**).

Next, differentially abundant bacterial taxa in the faecal samples at the phylum level were analysed between pairs of groups; healthy *vs.* cADpre, cADtreat *vs.* healthy, and cADpre *vs.* cADpost (**Fig. S9**, **Table 1**). In untreated cAD-affected (cADpre) dogs, the abundance of *Fusobacteria* was significantly lower than that in healthy dogs (effect size = −5.3705, *q* = 1.38×10^-16^), while the abundance was increased after 2 weeks of oclacitinib treatment (effect size = 5.11224, *q* < 0.0001). No change in this specific bacterial phylum was observed when cAD-affected dogs that already received oclacitinib treatment (cADtreat) and healthy dogs were compared.

**Table 1.** Statistical summary of the differentially abundant faecal bacterial taxa at the phylum level.

| Comparison | Phylum | fdr | Effect | Error min | Error max |
| --- | --- | --- | --- | --- | --- |
| cADpre vs. Healthy | *Fusobacteria* | 0.0000 | -5.3705 | -6.6151 | -4.1256 |
| cADpost vs. cADpre | *Actinobacteria* | 0.0296 | -0.4049 | -0.7696 | -0.0401 |
|  | *Bacteroidetes* | 0.0000 | -0.4057 | -0.4932 | -0.4201 |
|  | *Firmicutes* | 0.0000 | -0.9028 | -0.9374 | -0.8681 |
|  | *Fusobacteria* | 0.0000 | 5.1122 | 4.8750 | 5.3494 |
|  | *Proteobacteria* | 0.0000 | 0.3675 | 0.3186 | 0.4164 |
| cADtreatment vs. Healthy | *Campylobacterota* | 0.0059 | -5.4319 | -8.6639 | -2.1999 |

Bacterial taxa with its composition greater than 0.0001% (relative value > 0.00001) are displayed. 0.000 in the fdr column denotes fdr corrected *p* values below 0.001.

Similarly, differentially abundant faecal bacterial taxa between the groups at the genus level were analysed (**Fig. 4c** and **Table 2**). The abundance of *Fusobacterium* was significantly lower in untreated cAD (cADpre) dogs than in healthy dogs (effect size = −5.3391, *q* = 3.64×10^-10^). It was increased in the same cAD-affected dogs after they received 2-week-oclacitinib treatment (cADpost *vs.* cADpre; effect size = 5.1177, *q* < 0.0001). A similar trend (increase in cADpre and decrease after the 2-week treatment) was observed in *Blautia* (effect size = −1.1738, *q* = 3.47×10^-245^), *Clostridium sensu stricto* (effect size = −1.2619, *q* = 5.67×10^-58^) and *Escherichia/Shigella* (effect size = −0.4567, *q* = 8.7837×10^-106^). Dogs in the cADtreat group did not show a reduction in the abundance of *Blautia* and *Clostridium sensu stricto*, which remained higher than that in healthy dogs (*Blautia*, *q* = n.s; *C. sensu stricto*, effect size = 1.6453, *q* = 4.80×10^-28^).

**Table 2.** Statistical summary of the differentially abundant faecal bacterial taxa at the genus level.

| Comparison | Genus | fdr | Effect | Error min | Error max |
| --- | --- | --- | --- | --- | --- |
| cADpre vs. Healthy | *Clostridium_sensu_stricto* | 0.0009 | 2.9500 | 1.4118 | 4.4882 |
|  | *Staphylococcus* | 0.0000 | 5.6982 | 4.2895 | 7.1068 |
|  | *Amedibacterium* | 0.0164 | 2.7795 | 0.8068 | 4.7522 |
|  | *Blautia* | 0.0195 | 1.6524 | 0.4350 | 2.8698 |
|  | *Anaerostipes* | 0.0000 | 3.4001 | 2.0319 | 4.7682 |
|  | *Escherichia / Shigella* | 0.0164 | 2.4102 | 0.7242 | 4.0962 |
|  | *Flavonifractor* | 0.0041 | 2.3061 | 0.9298 | 3.6824 |
|  | *Fusobacterium* | 0.0000 | -5.2969 | -6.8852 | -3.7086 |
|  | *Porphyromonas* | 0.0284 | 5.8251 | 1.2393 | 10.4108 |
| cADpost vs. cADpre | *Bacteroides* | 0.0000 | -1.1320 | -1.1734 | -1.0906 |
|  | *Clostridium_sensu_stricto* | 0.0000 | -1.2757 | -1.4293 | -1.0906 |
|  | *Romboutsia* | 0.0000 | 0.8737 | 0.5212 | -1.1221 |
|  | *Terrisporobacter* | 0.0197 | -0.4578 | -0.8307 | -0.0848 |
|  | *Staphylococcus* | 0.0000 | 2.1513 | 1.9130 | 2.3895 |
|  | *Amedibacterium* | 0.0000 | -3.2703 | -3.4783 | -3.0623 |
|  | *Erysipelatoclostridium* | 0.0003 | -0.9847 | -1.5052 | -0.4642 |
|  | *Faecalimonas* | 0.0000 | -0.7794 | -0.8783 | -0.6806 |
|  | *Lachnospiraceae_incertae_sedis* | 0.0000 | 0.3221 | 0.2507 | 0.3936 |
|  | *Roseburia* | 0.0000 | 1.4039 | 1.1311 | 1.6767 |
|  | *Blautia* | 0.0000 | -1.1882 | -1.2569 | -1.1196 |
|  | *Escherichia / Shigella* | 0.0000 | -0.5029 | -0.5435 | -0.4624 |
|  | *Flavonifractor* | 0.0000 | -0.9294 | -1.0764 | -0.7824 |
|  | *Enterocloster* | 0.0002 | 2.5604 | 1.2616 | 3.8592 |
|  | *Anaerotignum* | 0.0000 | 1.4712 | 0.8140 | 2.1285 |
|  | *Fusobacterium* | 0.0000 | 5.0975 | 4.8608 | 5.3343 |
|  | *Proteus* | 0.0000 | 2.6491 | 2.5275 | 2.7707 |
|  | *Porphyromonas* | 0.0000 | 4.8906 | 4.5019 | 5.2792 |
| cADtreatment vs.<br>Healthy | *Clostridium_sensu_stricto* | 0.0000 | 1.5080 | 1.0417 | 1.9743 |
|  | *Turicibacter* | 0.0758 | 4.9954 | 1.0507 | 8.9401 |
|  | *Anaerotignum* | 0.0014 | -5.0315 | -7.6287 | -2.4342 |
|  | *Faecalimonas* | 0.0000 | -1.0094 | -1.1468 | -0.8719 |
|  | *Blautia* | 0.0889 | 1.8236 | 0.3075 | 3.3396 |
|  | *Corynebacterium* | 0.0758 | -2.9621 | -5.2842 | -0.6399 |

Bacterial taxa with its composition greater than 1% (relative value > 0.01) are presented. 0.000 in the *fdr* column denotes *fdr* corrected *p* values below 0.001.

### Correlation between faecal bacterial taxa and clinical cAD parameters

Next, we conducted a correlation analysis between clinical parameters (CADESI-04 total score and PVAS; **Fig. S10**) and bacterial taxa in the gut in each group. To evaluate specific bacterial phyla and/or genera correlating with respective clinical phenotypes commonly shared in all dogs, we looked for similar trends of correlation in all four dog groups. No bacterial phyla or other taxa were observed to correlate with PVAS (**Fig. S10a, b**). Three phyla (*Bacteroidetes*, *Actinobacteria* and *Firmicutes*) and three genera (*Amedibacillus*, *Bacteroides*, and *Flavonifractor*) were positively correlated with CADESI-04 total score, while three phyla (*Acidobacteria*, *Proteobacteria*, and *Synergistetes*) and four genera (*Cellulosilyticum*, *Escherichia/Shigella*, *Novosphingobium*, and *Stenotrophobacter*) were inversely correlated with the score (**Fig. S10c, d**).

### Differential gut microbial diversity in cAD-affected dogs compared with healthy dogs was reversed by oclacitinib treatment

Species diversity in the faecal samples was calculated using the DivNet estimate of Shannon that integrates the number of different species present (species richness) as well as their relative distribution (species evenness). DivNet also accounts for interactions between bacterial communities in the ecological network and for missing taxa. When alpha diversity was measured samplewise, no significant differences were observed between any of the four groups (**Fig. 4d**). Samples from all groups showed a wide range of species diversities inside their group, with mean values of 2.01 (healthy), 1.88 (cADpre), 1.96 (cADpost), and 2.06 (cADtreat). Therefore, there was a tendency towards a reduction in species diversity in untreated cAD-affected dogs (cADpre) compared with healthy dogs. Next, we conducted a communitywise estimate, which resulted in statistically significant differences between the groups (**Fig. 4e**). In this analysis, all samples per group are combined and regarded as one community so that the overall species diversity of the healthy or diseased environment is estimated. Compared with healthy dogs, community diversity was significantly decreased in untreated cAD (cADpre) dogs (healthy, 2.06 ± 0.35, *p* < 0.001; ADpre, 1.01 ± 0.73, *p* < 0.001). Similarly, the community after two weeks of treatment was less diverse compared to the healthy group (0.96 ± 0.57, *p* < 0.001). In the cAD groups that already received oclacitinib treatment (cADtreat), the mean diversity became closer to that of healthy dogs (1.92 ± 0.66), although a larger deviation between individuals was observed.

Beta diversity measures dissimilarity between the bacterial composition of multiple populations. The “Aitchison distance”, which is the Euclidian distance after *clr* (centred log-ratio) transformation, was used to measure beta diversity and constrained analysis of principal coordinates to visualize and evaluate clustering of the samples according to study group and disease status (**Fig. 4f**).

To analyse the ability of the samples to be separated by the group (Healthy, cADpre, cADpost and cADtreat) and disease severity (CADESI-04 score), multivariate analysis of variance using the distance matrices (adonis) was used. The disease separate categories were applied according to the published consensus [49], i.e., none, <10; mild cAD, 10-34; moderate cAD, 35-59, and severe cAD, ≥60. Aitchison distances showed that the group explained 14.6% (*p* < 0.0001) and disease severity explained 11.5% (*p* < 0.001) of the variance (*r^2^*) and that the healthy dogs were found to be significantly different compared with each of the other cAD groups (PERMANOVA, *p* < 0.001 for cADpre and cADpost; *p* < 0.001 for cADtreat). When testing for disease severity, beta diversity significantly differed between disease-free animals (none) and those with mild cAD (*p* < 0.05), while other comparisons did not reach statistical significance. Neither the sex (adonis, *p* = 0.64) of the dogs nor the age group (adonis, *p* = 0.09) was found to be significant in beta diversity. In summary, the bacterial community in all dogs with cAD, regardless of the treatment status, was significantly different from that in healthy dogs.

### Mitochondrial DNA variant clustering and mitochondrial DNA haplogroup determination

Buccal swab samples were obtained, and gDNA was isolated from all 40 dogs (20 healthy, 10 cADpre, and 10 cADtreat). Two swab samples did not yield sufficient gDNA for further processing. Ultimately, samples from 20 healthy dogs, nine cADpre dogs, and nine cADtreatment dogs were successfully sequenced for whole canine mitochondrial DNA. The obtained sequencing data were aligned to the reference sequence and variant calling was performed.

**Fig. S11** shows the complete mitochondrial genetic profiles of all 38 dogs. For haplogroup determination, we first used the Canis mtDNA HV1 database, which is available online (http://chd.vnbiology.com). The haplogroups determined by this approach overlapped the heatmap clustering pattern. Therefore, we decided to use major haplogroup A (combined all A sub-haplogroups, except A64; a total of 20 dogs, of which 16 had microbiome data available) and haplogroup C17 (including three samples assigned “new haplogroup C ref: C17”, thus together referred to as C afterwards; a total of 14 dogs, of which 12 dogs had microbiota data available) for further association analysis.

### Mitochondrial DNA haplogroups are associated with the gut microbiome

We hypothesized that genetic variation in the mtDNA is associated with cAD. Since all homoplasmic variants in the tested dogs are haplogroup-specific (**Fig. S11**), we performed a χ^2^ test for the mtDNA haplogroup distribution between healthy and diseased samples (cADpre and cADtreat). There was no association between the mtDNA haplogroup and cAD in this cohort (*p* = 0.2675).

Next, to evaluate the association between mtDNA haplogroup and alpha diversity, a random effects model (no random slope/intercept) was used with group (healthy, cADpost, and cADtreat) as a random effect (**Fig. 5a**). The model shows that the effect of the haplogroup on alpha diversity is not significant (*p* = 0.1194). Additionally, testing for heterogeneity of total diversity revealed a significant reduction in alpha diversity in haplogroup C (**Fig. 5b**; beta: *p* = 0.033, estimate = −0.3730).

**Fig. 5.**
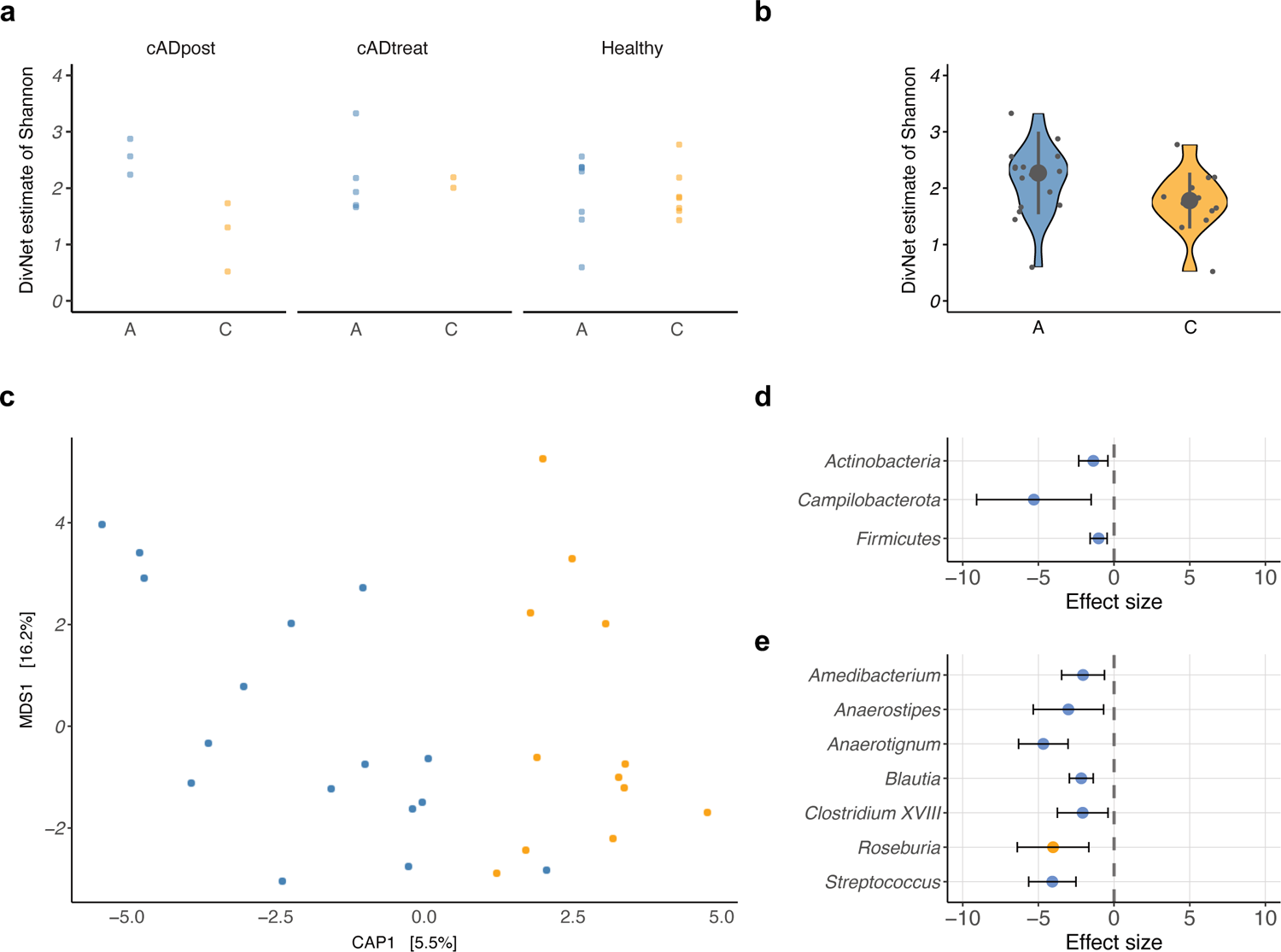
An association between the mitochondrial haplogroup and gut microbiota was evaluated. Alpha diversity was compared between faecal bacteria of dogs with haplogroup A and that of dogs with haplogroup C in 3 groups: cADpost, cADtreatment and healthy (**a**, left panel), and in all three groups together (**b**, right panel). When comparing haplogroups A and C in all three groups (regardless of health status), alpha diversity in dogs with haplogroup A showed a higher trend than that in dogs with haplogroup C (*p* = 0.074). Beta diversity was also compared between these two haplogroups (**c**). The Aitchison distance between haplogroups A and C demonstrated a difference with *p* = 0.078. The differentially abundant bacterial taxa between haplogroups A and C at the phylum level (**d**) and at the genus level (**e**) are presented (*q* < 0.1); blue dots refer to genera identified by corncob, and orange dots refer to genera identified by corncob and ANCOM-BC.

Beta diversity was estimated using Aitchison distance, and a constrained correspondence analysis was applied to estimate differences and correct for group effects (estimating the haplogroup effects; **Fig. 5c**). The model was not significant (*p* = 0.07822). Neither haplogroup (*p* = 0.07845) nor the separation at the x-axis (CAP1, *p* = 0.0784) were significant; *p* values were calculated using 99,999 permutations using an ANOVA-like permutation test for constrained correspondence analysis. Although these values were not statistically significant, they did show a trend (*p* < 0.1).

Finally, we analysed the differential abundance using corncob analysis (**Fig. 5d, e**). Three phyla (*q* < 0.01; *Actinobacteria*, *Campylobacterota*, and *Firmicutes*) and seven genera (*Amedibacterium*, *Anaerostipes*, *Anaerotignum*, *Blautia*, *Clostridium XVII*, *Roseburia*, and S*treptococcus*) were significantly lower in the C haplogroup than in the A haplogroup (*q* < 0.1). For this analysis, group was included in the null model to remove group effects from the analysis.

In summary, there was a trend of differences in beta diversity, and we observed significantly different abundant phyla/genera.

## Discussion

The skin and commensal microbiota are critical players in the health and pathological status of hosts. AD is an allergic skin disorder observed in both humans and dogs.

In this study, 16S rRNA gene amplicon sequencing of skin swab and stool samples obtained from dogs affected with cAD and healthy Shiba Inu dogs was conducted.

From the skin microbiota analysis, we demonstrated clear dysbiosis on the skin of cAD-affected dogs, with a higher abundance of *Staphylococcus* in all 12 skin sites of cAD-affected Shiba Inu dogs. These findings are in line with previously reported studies. The paired samples, i.e., untreated cAD (cADpre) and the same cAD-affected dogs after 2 weeks of oclacitinib treatment (cADpost), demonstrated that *Staphylococcus* abundance, predominantly *S. pseudintermedius*, was significantly lower in cADpost dog skin than in cADpre dog skin. Upon oclacitinib treatment, the abundance was significantly reduced.

To the best of our knowledge, no study has investigated the faecal microbiota in cAD to date. Additionally, we were able to evaluate the therapeutic impact of oclacitinib on the gut microbiota of untreated cAD-affected dogs that received 2 weeks or cAD dogs that were already on the treatment for an average of 13 months with oclacitinib. In fact, because untreated cAD-affected dogs (cADpre) and 2-week oclacitinib-treated cAD-affected dogs were paired, we were able to evaluate the gut microbiota in pre- and post-treatment cAD status. In parallel to the skin microbiota, there was clear dysbiosis in the gut of cAD-affected dogs. In healthy dogs, the predominant phyla in the faecal samples included *Firmicutes*, *Bacteroidetes*, *Proteobacteria* and *Fusobacteria*. This observation is in line with previous studies [50]. When comparing the gut microbiota in healthy dogs with that in cAD-affected dogs (cADpre), the abundance of *Fusobacteria* was significantly reduced in cAD-affected dogs. This was further illustrated at the genus level, presenting the reduction of *Fusobacterium* abundance in cADpre dogs compared with that in healthy dogs. In fact, *Fusobacterium* was primarily detected in several carnivores (African lion, arctic fox, king vulture) and non-carnivores (dogs and meerkat) as commensal microbiota [51]. In humans, *Fusobacterium* is known to be highly abundant in various gastrointestinal diseases, such as inflammatory bowel disease (IBD) [52], ulcerative colitis [53] and colorectal cancer [54], suggesting that the impact of *Fusobacterium* on health and disease in dogs seems to be different from that in humans [55].

*Megamonas*, which belongs to the phylum *Firmicutes* (*Bachillota*), is another bacterial genus that was significantly less abundant in cADpre dogs than in healthy dogs. These bacterial taxa are known to be pathologically relevant, such as in IBD in dogs; more specifically, the abundances of both *Fusobacterium* and *Megamonas* were negatively correlated with canine IBD [56]. In dogs, the abundance of *Fusobacterium* is positively correlated with faecal concentrations of butyrate and propionate [57], and some species of *Megamonas* can ferment glucose into acetate and propionate [58, 59]. Given that the higher levels of butyrate and propionate in faeces reduced the risk of atopy in early life in humans, the reduction of such SCFA-producing bacteria in cAD-affected dogs suggests that *Fusobacteria* and *Megamonas* could explain the pathological relevance of cAD in Shiba Inu dogs. Furthermore, several meta-analysis studies of the published data suggested an association between IBD and AD in humans, i.e., a higher incidence of AD in IBD patients as well as a higher prevalence of IBD in AD patients [60, 61], suggesting that the phylum *Fusobacteria* and the genus *Fusobacterium* may be potential biomarkers not only for the risk of cAD but also for the risk of IBD in dogs. Nevertheless, further studies of faecal microbiota from many different dog breeds will be needed to confirm this hypothesis.

In contrast, the abundances of *Clostridium sensu stricto* (former *Clostridium cluster I*) and *Escherichia/Shigella* were significantly increased in untreated cAD-affected dogs (cADpre) compared with healthy dogs. *Clostridium* was reported that its colonization in the gut at an early age (at ages 5 and 13 weeks) was associated with a higher risk of AD in children [62] and that the higher abundance of *Clostridium spp*. is associated with AD in infants. Two bacterial taxa, *Escherichia coli* and *Shigella spp.*, were not differentiated by the 16S rRNA gene sequence because of their genetic relatedness [63, 64]. Despite some differential biochemical characteristics, both bacterial taxa are enteric pathogens, and their virulence factors influence many cellular processes [64]. Higher abundances of both *Shigella* and *E. coli* in faeces were reported to be associated with atopic eczema [65, 66].

These changes observed in the gut microbiota of the untreated cAD-affected dogs (cADpre) were further altered most likely by medical intervention with oclacitinib. After the 2-week treatment, the abundance of *Fusobacterium*, which was less than 2% in cAD-affected dogs, was significantly increased in the same dogs (cADpost *vs.* cADpre). On the other hand, the abundance of *Escherichia/Shigella* was significantly decreased after the 2-week treatment. In cAD-affected dogs that were already on oclacitinib treatment (cADtreat), the composition of the gut bacterial taxa was richer than that of the cAD-affected dogs that received 2-week treatment (cADpost): *Fusobacterium* was retained, the abundance of *Escherichia/Shigella* was reduced, and *Megamonas* appeared. This finding indicates that the 2-week oclacitinib treatment shifted gut microbial composition, including the increased abundance of *Fusobacterium*, which may be beneficial for dog health. The gut microbial composition of the faeces from cAD-affected dogs that were already on oclacitinib treatment (cADtreat) showed relatively similar patterns compared with untreated cAD-affected dogs (cADpre), i.e., higher abundances of *Escherichia/Shigella* and *Clostridium sensu stricto*, yet we observed higher abundances of *Fusobacterium* and *Megamonas*, which were also highly abundant in healthy dogs, suggesting that the treatment with oclacitinib may shift the gut microbial composition of cAD-affected dogs towards that of healthy dogs to a certain extent. Five out of 10 cADtreat dogs that already received oclacitinib treatment, were on a varied prescription diet specifically designed for allergy control; thus, this dietary control may have also supported modulating the gut microbiota composition from “cAD gut type” to “healthy gut type”.

We also sequenced the whole mitochondrial genome of Shiba Inu dogs with and without cAD in this study. In humans, mitochondrial haplogroups are utilized in the context of population genetics because they represent a phylogenetic tree that allows retracing human settlement of the world, resulting in present-day worldwide geographic differences in haplogroup abundances. In contrast, dogs have been recently crossed, and their breeding has been largely controlled by humans. Therefore, we hypothesized that mtDNA variants may be variable among one specific breed, which may contribute to host gene-microbiota interactions in the pathology of cAD. We did not observe a significant association between mtDNA haplogroups and cAD in this sample cohort; however, this may be due to the limited sample size. Yet we identified a significant association between mtDNA haplogroups and specific bacterial taxa. Further studies using larger sample sizes may indicate that the mtDNA haplogroup and specific gut microbes are biomarkers to evaluate disease predisposition, treatment efficacy, and health management in dogs.

## Conclusions

We conducted a comprehensive microbial analysis in the skin and gut of cAD and healthy Shiba Inu dogs. The cAD-affected dogs included untreated cAD-affected dogs, the same dogs after a 2-week course of oclacitinib treatment, and an independent group of cAD-affected dogs that had already been receiving oclacitinib treatment for up to two years. This study is the first report to present clear dysbiosis of both the skin and the gut in dogs with cAD. We observed that oclacitinib treatment on cAD, regardless the treatment duration, shifted both the skin and gut microbiota toward those of healthy dogs. In addition, we identified a significant association between mtDNA haplogroups and specific bacterial taxa in the gut microbiota in dogs. This finding will be the basis for novel disease management strategies for canine cAD.

## Availability of data and materials

Skin and faecal 16S rRNA sequencing data and dog mitochondrial genome sequencing data were submitted to the European Nucleotide Archive (ENA) and are available under accession numbers PRJEB53048 (microbiome data) and PRJEB53275 (whole mitochondrial genome sequencing data).

## List of abbreviations

AD: atopic dermatitis

cAD: canine atopic dermatitis

CADESI-04: canine atopic dermatitis evaluation score index - 04

PVAS: pruritis visual analogue scale

16SrRNA: 16S ribosomal RNA.

## Acknowledgements

We thank the dogs and their owners for participating in this study. Authors also thank Ms. Miriam Freitag and Ms. Petra Langenstrassen for their excellent technical assistance. We acknowledge to Dr. Atsushi Yamamoto for supporting communications and for organising clinical data and samples. MT, AK, IW, MO, TL and HB acknowledge computational support from the OMICS compute cluster at the University of Lübeck. IW and HB acknowledge funding by the Deutsche Forschungsgemeinschaft (DFG, German Research Foundation) under Germany’s Excellence Strategy - EXC 22167-390884018. We would like to thank the late Dr. Ayako Kamida for supporting clinical samples organisation at the initial phase of the project.

## Author information

### Authors’ contributions

MT, AK and MO conducted bioinformatics analysis for skin and gut microbiota. IW and TL performed bioinformatics analysis for mitochondrial DNA. KN, TO and YS made the clinical diagnosis and collected skin swab samples, stool samples and buccal swab samples from dogs. AW was involved in the project conceptualisation, coordinated the recruitment and logistics of the samples, and contributed to the data interpretation. SMI and HB were involved in the project conceptualisation and data interpretation. MH performed experiments, designed and directed the study. MT, AK, IW and MH drafted the manuscript, and all authors contributed to finalize the manuscript.

### Corresponding authors

Correspondence to Misa Hirose.

### Ethics declarations

Ethics approval and consent to participate, The Clinical Animal Trials and Research Ethics Committee of the Tokyo University of Agriculture and Technology approved all studies (approval no. 0016021). All participating dog owners were provided with information about the study and gave their written consent before their dogs were included.

### Consent for publication

All authors have reviewed the manuscript.

## Supplementary figures

**Fig. S1** The cohort of Shiba Inu dogs tested in this study (**a**). Twelve skin swab samples and a faecal sample were collected from each dog in each group, resulted in a total of 12 × (10 + 10+ 10 + 20) = 600 skin swab samples and 1 × (10 + 10 + 10 + 20) = 50 faecal samples. A buccal sample was collected from three groups; cADpre, cADtreat and Healthy, 1 × (10 + 10 + 20) = 40 samples. Age distribution of the participating dogs (**b**), CADESI-04 (**b**) and PVAS (**c**) values for each dog were plotted in the graphs. The values of healthy dogs were given as 0. Ten dogs with untreated cAD dogs (cADpre) received oclacitinib treatment for 2 weeks (cADpost). The treatment efficacy was shown as a reduction in CADESI-04 (**d**) and PVAS values (**e**). For these 10 dogs, the CADESI-04 score in 12 different skin sites, the perilabial area, medial pinna, axilla, front paw, hind paw, cubital flexor, palmar metacarpal, flank, inguinal area, abdomen, perineum, and ventral tail, is shown.

**Fig. S2** For alpha diversity, Shannon indices were compared between four groups: healthy dogs (Healthy), cAD-affected dogs without medical treatment (cADpre), cADpre dogs that received 2 weeks of oclacitinib treatment (cADpost), and cAD-affected dogs already on oclacitinib treatment (cADtreat). The comparisons were made in four selected skin sites: abdomen (**a**), front paw (**b**), perilabial area (**c**), and ventral tail (**d**). *P* values showing significant correlations (*p* < 0.05) are presented.

**Fig. S3** Beta diversity was analysed by CAPscale analysis of Aichison distances and was compared between four groups: healthy dogs (Healthy), cAD-affected dogs without medical treatment (cADpre), cADpre dogs that received 2 weeks of oclacitinib treatment (cADpost), and cAD-affected dogs already on oclacitinib treatment (cADtreat) in four selected skin sites: abdomen (**a**), front paw (**b**), perilabial area (**c**), and ventral tail (**d**).

**Fig. S4** Differentially abundant bacterial taxa at the phylum level between two groups; healthy and cADpre, cADtreat and healthy, or cADpre and cADpost in four selected skin sites; abdomen (**a**), front paw (**b**), perilabial area (**c**), and ventral tail (**d**). Differentially abundant taxa (*q* < 0.1) are presented in the plots; blue dots refer to phyla identified by corncob, and orange dots refer to phyla identified by corncob and ANCOM-BC.

**Fig. S5** Differentially abundant bacterial taxa at the genus level between two groups; healthy and cADpre, cADtreat and healthy, or cADpre and cADpost in four selected skin sites; abdomen (**a**), front paw (**b**), perilabial area (**c**), and ventral tail (**d**). Differentially abundant taxa (*q* < 0.1) are presented in the plots; blue dots refer to genera identified by corncob, and orange dots refer to genera identified by corncob and ANCOM-BC.

**Fig. S6** Correlation of pruritis score (PVAS) and bacterial taxa at the phylum level (**a**) and at the genus level (**b**) in the four skin sites: abdomen, front paw, perilabial area and ventral tail. *P* values showing significant correlations (*p* < 0.05) are presented.

**Fig. S7** Of the twelve investigated skin sites, the abundance of *Staphylococcus* was associated with cAD in the skin sites, showing a significant difference at five sites when compared between dogs with untreated cAD (cADpre) and healthy dogs; these sites were the abdomen (**a**), axilla (**b**), cubital flexor (**c**), palmar metacarpal (**d**), and perilabial area (**e**). Three sites, the flank (**f**), front paw (**g**), and hind paw (**h**), also showed a tendency towards a higher abundance of *Staphylococcus* in cADpre dogs than in healthy dogs. Marginal effects for each comparison are shown in the left column plots with 95% confidence intervals (whiskers); the *p* value for the comparison between cADpre and Healthy is shown. The *p* values were corrected using the Tukey method for comparing a family of three estimates. The abundance of *Staphylococcus* in each of these skin sites correlated with its respective CADESI-04 score (**i**-**p**).

**Fig. S8** Data related to Fig. 3. The mean abundances of the 10 predominant *Staphylococcus* spp. are presented in the four skin sites: abdomen, front paw, perilabial area, and ventral tail. Abundances are shown per study group; healthy dogs, cAD-affected dogs before treatment (cADpre), cADpre dogs that received 2 weeks of oclacitinib treatment (cADpost) and cAD-affected dogs already on treatment (cADtreat) are presented.

**Fig. S9** Data related to Fig. 4. Differentially abundant taxa in the gut at the phylum level between the two groups (*p* < 0.1) are shown. The groups compared are indicated in each panel: (**left**) cAD-affected dogs that did not receive medical treatment (cADpre) *vs.* healthy dogs (Healthy); (**middle**) cAD-affected dogs on medical treatment (cADtreat) *vs.* Healthy; (**right**) cADpre dogs that received 2-week oclacitinib treatment (cADpost) *vs.* cADpre. Blue dots refer to phyla identified by corncob, and orange dots refer to phyla identified by corncob and ANCOM-BC.

**Fig. S10** Correlation of bacterial taxa in the gut and canine clinical score (CADESI-04; total score; **a** and **b**) or pruritis score (PVAS; **c** and **d**) at the phylum level (**a** and **c**) and at the genus level (**b** and **d**). *P* values showing significant correlations (*p* < 0.05) are presented.

**Fig. S11** The PHRED-scaled genotype likelihood of nucleotide variants in the mitochondrial genome identified in each dog are presented in a heatmap. This value ranges between 0 and 255, with 0 being the strongest support of a variant (displayed in black). Positions with no variant call are displayed in white. These variant patterns divide the dogs into three clusters which agree with the mitochondrial haplogroups assigned: A, B and C. The two mitochondrial haplogroups with more than 10 dogs included, haplogroup A (excluding A64, because its variant profile is markedly different) and C, were selected for further analysis of the faecal microbiota.

## Supplementary tables (Supplementary_file_TableS1-S6.xlsx)

**Table S1.** Age and sex distribution of dogs involved in the study.

**Table S2.** Correlation analysis between bacterial taxa and clinical score (CADESI-04) to select the skin sites for further analysis.

**Table S3.** Correlation of *Staphylococcus* abundance and CADESI-04 score in each skin site was compared between dog groups.

**Table S4.** Mean relative abundance of the 10 most abundant *Staphylococcus spp.* in the selected four skin samples form healthy, cADpre, cADpost and cADtreat dogs. Percent identity of the best hit from NCBI Nucleotide BLAST is presented in each classified species.

**Table S5.** Mean relative abundance of bacterial taxa at the phylum level in faecal samples from all four dog groups.

**Table S6.** Mean relative abundance of bacterial taxa at the genus level in the faecal samples in all four dog groups.

## References

1. Hayward JJ, Castelhano MG, Oliveira KC, Corey E, Balkman C, Baxter TL, et al. Complex disease and phenotype mapping in the domestic dog. Nat Commun. 2016;7:10460.

2. LeBlanc AK, Mazcko CN. Improving human cancer therapy through the evaluation of pet dogs. Nat Rev Cancer. Nature Publishing Group; 2020;20:727–42.

3. Ruple A, MacLean E, Snyder-Mackler N, Creevy KE, Promislow D. Dog Models of Aging. Annu Rev Anim Biosci. 2022;10:419–39.

4. Weidinger S, Beck LA, Bieber T, Kabashima K, Irvine AD. Atopic dermatitis. Nat Rev Dis Primer. Nature Publishing Group; 2018;4:1–20.

5. Gedon NKY, Mueller RS. Atopic dermatitis in cats and dogs: a difficult disease for animals and owners. Clin Transl Allergy. 2018;8:41.

6. Nuttall TJ, Marsella R, Rosenbaum MR, Gonzales AJ, Fadok VA. Update on pathogenesis, diagnosis, and treatment of atopic dermatitis in dogs. J Am Vet Med Assoc. 2019;254:1291– 300.

7. Marsella R, Girolomoni G. Canine models of atopic dermatitis: a useful tool with untapped potential. J Invest Dermatol. 2009;129:2351–7.

8. Jaeger K, Linek M, Power HT, Bettenay SV, Zabel S, Rosychuk R a. W, et al. Breed and site predispositions of dogs with atopic dermatitis: a comparison of five locations in three continents. Vet Dermatol. 2010;21:118–22.

9. Williams MR, Gallo RL. The role of the skin microbiome in atopic dermatitis. Curr Allergy Asthma Rep. 2015;15:65.

10. Carding S, Verbeke K, Vipond DT, Corfe BM, Owen LJ. Dysbiosis of the gut microbiota in disease. Microb Ecol Health Dis. 2015;26:26191.

11. Kong HH, Oh J, Deming C, Conlan S, Grice EA, Beatson MA, et al. Temporal shifts in the skin microbiome associated with disease flares and treatment in children with atopic dermatitis. Genome Res. 2012;22:850–9.

12. Rodrigues Hoffmann A, Patterson AP, Diesel A, Lawhon SD, Ly HJ, Elkins Stephenson C, et al. The skin microbiome in healthy and allergic dogs. PloS One. 2014;9:e83197.

13. Bradley CW, Morris DO, Rankin SC, Cain CL, Misic AM, Houser T, et al. Longitudinal Evaluation of the Skin Microbiome and Association with Microenvironment and Treatment in Canine Atopic Dermatitis. J Invest Dermatol. 2016;136:1182–90.

14. Chermprapai S, Ederveen THA, Broere F, Broens EM, Schlotter YM, van Schalkwijk S, et al. The bacterial and fungal microbiome of the skin of healthy dogs and dogs with atopic dermatitis and the impact of topical antimicrobial therapy, an exploratory study. Vet Microbiol. 2019;229:90–9.

15. Fazakerley J, Nuttall T, Sales D, Schmidt V, Carter SD, Hart CA, et al. Staphylococcal colonization of mucosal and lesional skin sites in atopic and healthy dogs. Vet Dermatol. 2009;20:179–84.

16. Lee SY, Lee E, Park YM, Hong SJ. Microbiome in the Gut-Skin Axis in Atopic Dermatitis. Allergy Asthma Immunol Res. 2018;10:354–62.

17. O’Neill CA, Monteleone G, McLaughlin JT, Paus R. The gut-skin axis in health and disease: A paradigm with therapeutic implications. BioEssays News Rev Mol Cell Dev Biol. 2016;38:1167–76.

18. Smith PM, Howitt MR, Panikov N, Michaud M, Gallini CA, Bohlooly-Y M, et al. The microbial metabolites, short-chain fatty acids, regulate colonic Treg cell homeostasis. Science. 2013;341:569–73.

19. Schwarz A, Bruhs A, Schwarz T. The Short-Chain Fatty Acid Sodium Butyrate Functions as a Regulator of the Skin Immune System. J Invest Dermatol. 2017;137:855–64.

20. Fang Z, Li L, Zhang H, Zhao J, Lu W, Chen W. Gut Microbiota, Probiotics, and Their Interactions in Prevention and Treatment of Atopic Dermatitis: A Review. Front Immunol [Internet]. 2021 [cited 2022 May 14];12. Available from: https://www.frontiersin.org/article/10.3389/fimmu.2021.720393

21. Tanaka K, Yamamoto-Fukuda M, Takizawa T, Shimakura H, Sakaguchi M. Association analysis of non-synonymous polymorphisms of interleukin-4 receptor-α and interleukin-13 genes in canine atopic dermatitis. J Vet Med Sci. 2020;82:1253–9.

22. Fujimura M, Ishimaru H, Nakatsuji Y. Fluoxetine (SSRI) treatment of canine atopic dermatitis: a randomized, double-blind, placebo-controlled, crossover trial. Pol J Vet Sci. 2014;17:371–3.

23. Wood SH, Ke X, Nuttall T, McEwan N, Ollier WE, Carter SD. Genome-wide association analysis of canine atopic dermatitis and identification of disease related SNPs. Immunogenetics. 2009;61:765–72.

24. Ma J, Coarfa C, Qin X, Bonnen PE, Milosavljevic A, Versalovic J, et al. mtDNA haplogroup and single nucleotide polymorphisms structure human microbiome communities. BMC Genomics. 2014;15:257.

25. Hirose M, Künstner A, Schilf P, Sünderhauf A, Rupp J, Jöhren O, et al. Mitochondrial gene polymorphism is associated with gut microbial communities in mice. Sci Rep. 2017;7:15293.

26. Hensel P, Santoro D, Favrot C, Hill P, Griffin C. Canine atopic dermatitis: detailed guidelines for diagnosis and allergen identification. BMC Vet Res. 2015;11:196.

27. Olivry T, Marsella R, Iwasaki T, Mueller R, Dermatitis TITFOCA. Validation of CADESI-03, a severity scale for clinical trials enrolling dogs with atopic dermatitis. Vet Dermatol. 2007;18:78–86.

28. Reimer-Taschenbrecker A, Künstner A, Hirose M, Hübner S, Gewert S, Ibrahim S, et al. Predominance of Staphylococcus correlates with wound burden and disease activity in dystrophic epidermolysis bullosa: a prospective case-control study. J Invest Dermatol. 2022;S0022–202X(22)00090-2.

29. Künstner A, Schilf P, Busch H, Ibrahim SM, Hirose M. Changes of Gut Microbiota by Natural mtDNA Variant Differences Augment Susceptibility to Metabolic Disease and Ageing. Int J Mol Sci. 2022;23:1056.

30. Human mtDNA Genome For the Illumina Sequencing Platform [Internet]. [cited 2019 Jan 30]. Available from: http://emea.support.illumina.com/content/dam/illumina-support/documents/documentation/chemistry_documentation/samplepreps_legacy/human-mtdna-genome-guide-15037958-01.pdf

31. Rognes T, Flouri T, Nichols B, Quince C, Mahé F. VSEARCH: a versatile open source tool for metagenomics. PeerJ. 2016;4:e2584.

32. Edgar RC. UPARSE: highly accurate OTU sequences from microbial amplicon reads. Nat Methods. 2013;10:996–8.

33. Edgar RC. SINTAX: a simple non-Bayesian taxonomy classifier for 16S and ITS sequences [Internet]. bioRxiv; 2016 [cited 2022 May 31]. p. 074161. Available from: https://www.biorxiv.org/content/10.1101/074161v1

34. Cole JR, Wang Q, Fish JA, Chai B, McGarrell DM, Sun Y, et al. Ribosomal Database Project: data and tools for high throughput rRNA analysis. Nucleic Acids Res. 2014;42:D633–642.

35. Schloss PD, Westcott SL, Ryabin T, Hall JR, Hartmann M, Hollister EB, et al. Introducing mothur: open-source, platform-independent, community-supported software for describing and comparing microbial communities. Appl Environ Microbiol. 2009;75:7537–41.

36. Quast C, Pruesse E, Yilmaz P, Gerken J, Schweer T, Yarza P, et al. The SILVA ribosomal RNA gene database project: improved data processing and web-based tools. Nucleic Acids Res. 2013;41:D590–6.

37. Price MN, Dehal PS, Arkin AP. FastTree 2--approximately maximum-likelihood trees for large alignments. PloS One. 2010;5:e9490.

38. McMurdie PJ, Holmes S. phyloseq: An R Package for Reproducible Interactive Analysis and Graphics of Microbiome Census Data. PLOS ONE. 2013;8:e61217.

39. Davis NM, Proctor DM, Holmes SP, Relman DA, Callahan BJ. Simple statistical identification and removal of contaminant sequences in marker-gene and metagenomics data. Microbiome. 2018;6:226.

40. Willis AD, Martin BD. Estimating diversity in networked ecological communities. Biostat Oxf Engl. 2020;kxaa015.

41. Willis A, Bunge J, Whitman T. Improved detection of changes in species richness in high diversity microbial communities. J R Stat Soc Ser C Appl Stat. [Wiley, Royal Statistical Society]; 2017;66:963–77.

42. Aitchison J. The Statistical Analysis of Compositional Data. J R Stat Soc Ser B Methodol. [Royal Statistical Society, Wiley]; 1982;44:139–77.

43. Martin BD, Witten D, Willis AD. MODELING MICROBIAL ABUNDANCES AND DYSBIOSIS WITH BETA-BINOMIAL REGRESSION. Ann Appl Stat. 2020;14:94–115.

44. Lin H, Peddada SD. Analysis of compositions of microbiomes with bias correction. Nat Commun. 2020;11:3514.

45. Susin A, Wang Y, Lê Cao K-A, Calle ML. Variable selection in microbiome compositional data analysis. NAR Genomics Bioinforma. 2020;2:lqaa029.

46. Li H, Durbin R. Fast and accurate short read alignment with Burrows-Wheeler transform. Bioinforma Oxf Engl. 2009;25:1754–60.

47. Danecek P, Bonfield JK, Liddle J, Marshall J, Ohan V, Pollard MO, et al. Twelve years of SAMtools and BCFtools. GigaScience. 2021;10:giab008.

48. Thai QK, Chung DA, Tran H-D. Canis mtDNA HV1 database: a web-based tool for collecting and surveying Canis mtDNA HV1 haplotype in public database. BMC Genet. 2017;18:60.

49. Olivry T, Saridomichelakis M, Nuttall T, Bensignor E, Griffin CE, Hill PB, et al. Validation of the Canine Atopic Dermatitis Extent and Severity Index (CADESI)-4, a simplified severity scale for assessing skin lesions of atopic dermatitis in dogs. Vet Dermatol. 2014;25:77–85, e25.

50. Suchodolski JS, Camacho J, Steiner JM. Analysis of bacterial diversity in the canine duodenum, jejunum, ileum, and colon by comparative 16S rRNA gene analysis. FEMS Microbiol Ecol. 2008;66:567–78.

51. Vital M, Gao J, Rizzo M, Harrison T, Tiedje JM. Diet is a major factor governing the fecal butyrate-producing community structure across Mammalia, Aves and Reptilia. ISME J. 2015;9:832–43.

52. Strauss J, Kaplan GG, Beck PL, Rioux K, Panaccione R, Devinney R, et al. Invasive potential of gut mucosa-derived Fusobacterium nucleatum positively correlates with IBD status of the host. Inflamm Bowel Dis. 2011;17:1971–8.

53. Ohkusa T, Sato N, Ogihara T, Morita K, Ogawa M, Okayasu I. Fusobacterium varium localized in the colonic mucosa of patients with ulcerative colitis stimulates species-specific antibody. J Gastroenterol Hepatol. 2002;17:849–53.

54. Bullman S, Pedamallu CS, Sicinska E, Clancy TE, Zhang X, Cai D, et al. Analysis of Fusobacterium persistence and antibiotic response in colorectal cancer. Science. 2017;358:1443–8.

55. Vázquez-Baeza Y, Hyde ER, Suchodolski JS, Knight R. Dog and human inflammatory bowel disease rely on overlapping yet distinct dysbiosis networks. Nat Microbiol. 2016;1:16177.

56. Suchodolski JS, Dowd SE, Wilke V, Steiner JM, Jergens AE. 16S rRNA gene pyrosequencing reveals bacterial dysbiosis in the duodenum of dogs with idiopathic inflammatory bowel disease. PloS One. 2012;7:e39333.

57. Minamoto Y, Minamoto T, Isaiah A, Sattasathuchana P, Buono A, Rangachari VR, et al. Fecal short-chain fatty acid concentrations and dysbiosis in dogs with chronic enteropathy. J Vet Intern Med. 2019;33:1608–18.

58. Chevrot R, Carlotti A, Sopena V, Marchand P, Rosenfeld E. Megamonas rupellensis sp. nov., an anaerobe isolated from the caecum of a duck. Int J Syst Evol Microbiol. 2008;58:2921–4.

59. Sakon H, Nagai F, Morotomi M, Tanaka R. Sutterella parvirubra sp. nov. and Megamonas funiformis sp. nov., isolated from human faeces. Int J Syst Evol Microbiol. 2008;58:970–5.

60. Lee H, Lee JH, Koh S-J, Park H. Bidirectional relationship between atopic dermatitis and inflammatory bowel disease: A systematic review and meta-analysis. J Am Acad Dermatol. 2020;83:1385–94.

61. Shi X, Chen Q, Wang F. The Bidirectional Association between Inflammatory Bowel Disease and Atopic Dermatitis: A Systematic Review and Meta-Analysis. Dermatol Basel Switz. 2020;236:546–53.

62. Penders J, Gerhold K, Stobberingh EE, Thijs C, Zimmermann K, Lau S, et al. Establishment of the intestinal microbiota and its role for atopic dermatitis in early childhood. J Allergy Clin Immunol. 2013;132:601–607.e8.

63. Brenner DJ, Fanning GR, Steigerwalt AG, Orskov I, Orskov F. Polynucleotide sequence relatedness among three groups of pathogenic Escherichia coli strains. Infect Immun. 1972;6:308–15.

64. Devanga Ragupathi NK, Muthuirulandi Sethuvel DP, Inbanathan FY, Veeraraghavan B. Accurate differentiation of Escherichia coli and Shigella serogroups: challenges and strategies. New Microbes New Infect. 2018;21:58–62.

65. Hong P-Y, Lee BW, Aw M, Shek LPC, Yap GC, Chua KY, et al. Comparative analysis of fecal microbiota in infants with and without eczema. PloS One. 2010;5:e9964.

66. Penders J, Thijs C, van den Brandt PA, Kummeling I, Snijders B, Stelma F, et al. Gut microbiota composition and development of atopic manifestations in infancy: the KOALA Birth Cohort Study. Gut. 2007;56:661–7.

