## Supplementary material S1, S2, supplementary Table S1-S6 for "A comprehensive analysis of gut and skin microbiota in canine atopic dermatitis in Shiba Inu dogs": 20220711_FigS1-S11.pdf

#### Fig. S1

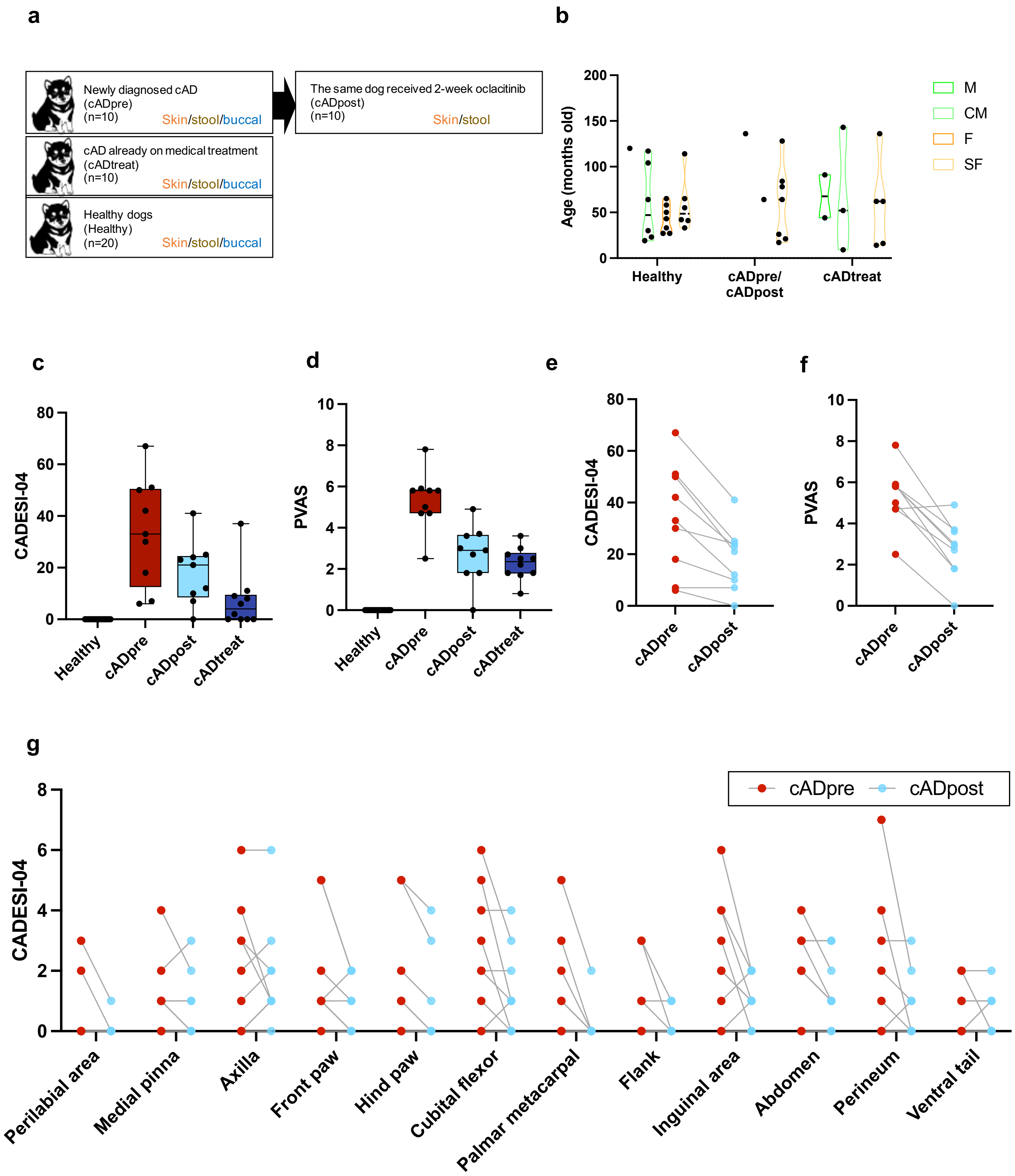

a
Newly diagnosed cAD
(cADpre)
(n=10)
Skin/stool/buccal
The same dog received 2-week oclacitinib
(cADpost)
(n=10)
Skin/stool
cAD already on medical treatment
(cADtreat)
(n=10)
Skin/stool/buccal
Healthy dogs
(Healthy)
(n=20)
Skin/stool/buccal
b
Age (months old)
Healthy
cADpre/
cADpost
cADtreat
M
CM
F
SF
c
CADESI-04
Healthy
cADpre
cADpost
cADtreat
d
PVAS
e
f
g
cADpre
cADpost
Perilabial area
Medial pinna
Axilla
Front paw
Hind paw
Cubital flexor
Palmar metacarpal
Flank
Inguinal area
Abdomen
Perineum
Ventral tail

**Fig. S2**

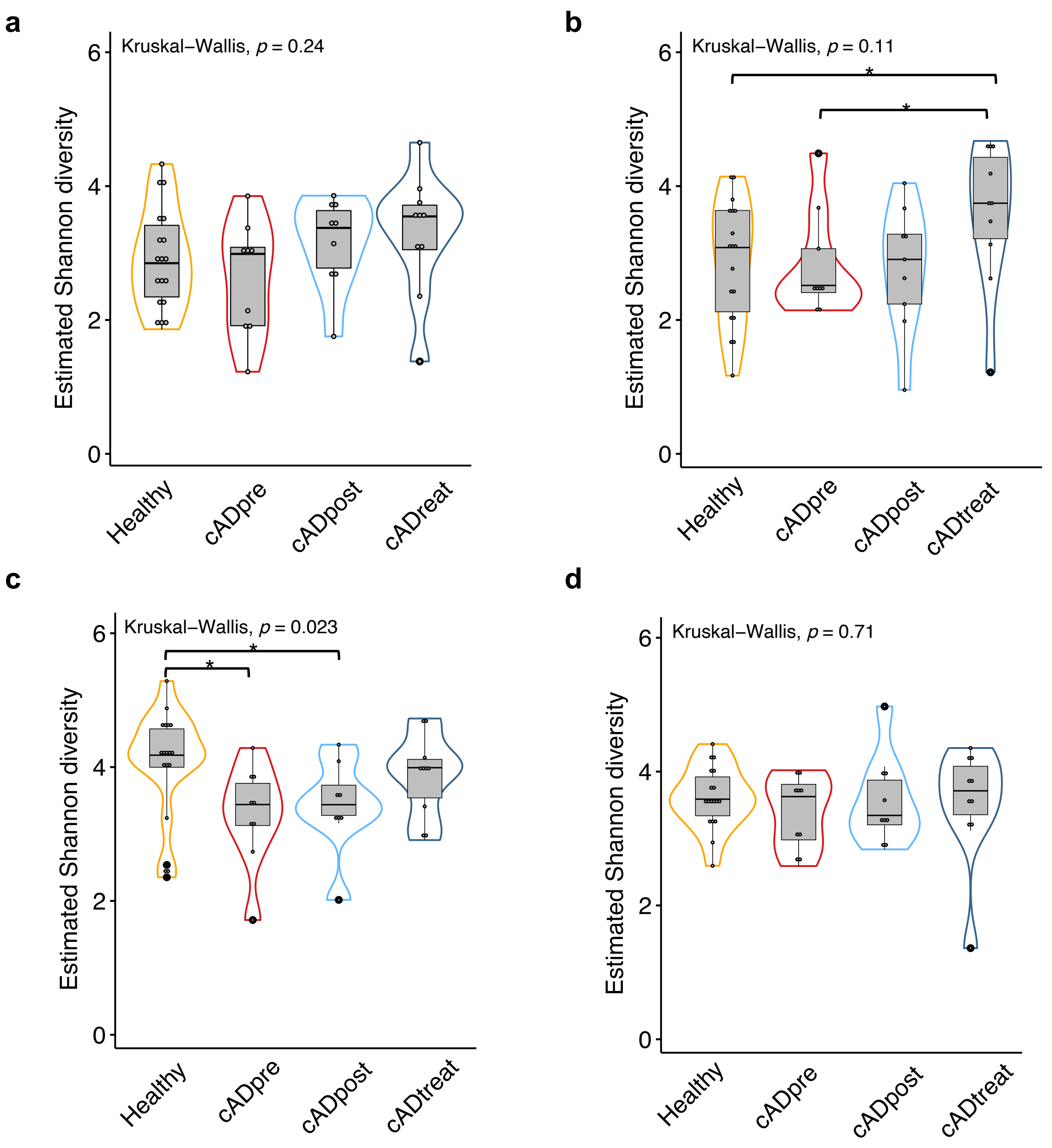
a
Kruskal–Wallis, p = 0.24
Estimated Shannon diversity
0
2
4
6
Healthy
cADpre
cADpost
cADreat
b
Kruskal–Wallis, p = 0.11
*
*
Estimated Shannon diversity
0
2
4
6
Healthy
cADpre
cADpost
cADtreat
c
Kruskal–Wallis, p = 0.023
*
*
Estimated Shannon diversity
0
2
4
6
Healthy
cADpre
cADpost
cADtreat
d
Kruskal–Wallis, p = 0.71
Estimated Shannon diversity
0
2
4
6
Healthy
cADpre
cADpost
cADtreat

**Fig. S3**

**a**

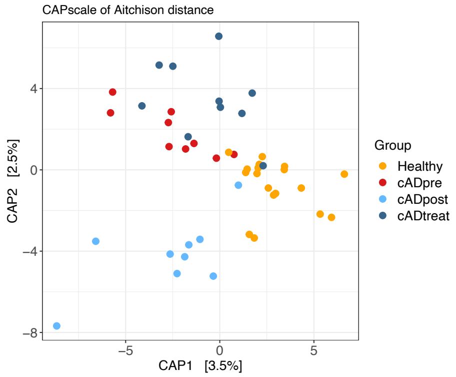


| | Df | Variance | F-statistic | p-value |
|---|---|---|---|---|
| Model | 3 | 210.55 | 1.2265 | 0.01073* |
| Residual | 42 | 2403.25 | | |

| | Df | Variance | F-statistic | p-value |
|---|---|---|---|---|
| Group | 3 | 210.55 | 1.2265 | 0.01129* |
| Residual | 42 | 2403.25 | | |

| | Df | Variance | F-statistic | p-value |
|---|---|---|---|---|
| CAP1 | 1 | 83.67 | 1.4623 | 0.0979 |
| Residual | 42 | 2403.25 | | |

**b**

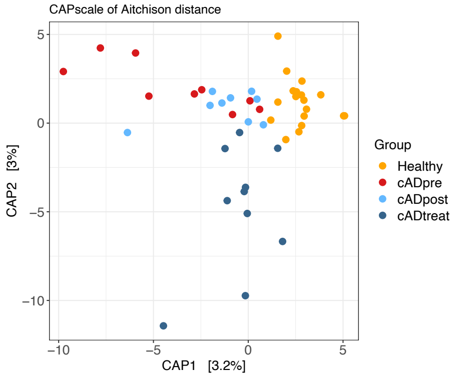


| | Df | Variance | F-statistic | p-value |
|---|---|---|---|---|
| Model | 3 | 211.47 | 1.2176 | 0.00063* |
| Residual | 42 | 2431.55 | | |

| | Df | Variance | F-statistic | p-value |
|---|---|---|---|---|
| Group | 3 | 211.47 | 1.2176 | 0.00063* |
| Residual | 42 | 2431.55 | | |

| | Df | Variance | F-statistic | p-value |
|---|---|---|---|---|
| CAP1 | 1 | 83.99 | 1.4507 | 0.02694* |
| CAP2 | 1 | 78.07 | 1.3484 | 0.03883* |
| Residual | 42 | 2431.55 | | |

**c**

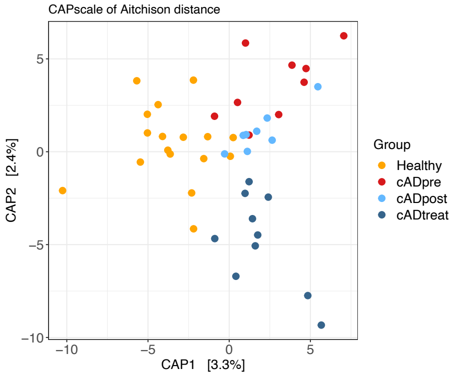


| | Df | Variance | F-statistic | p-value |
|---|---|---|---|---|
| Model | 3 | 292.7 | 1.089 | 0.1025 |
| Residual | 40 | 3584.1 | | |

| | Df | Variance | F-statistic | p-value |
|---|---|---|---|---|
| Group | 3 | 292.7 | 1.089 | 0.1027 |
| Residual | 40 | 3584.1 | | |

| | Df | Variance | F-statistic | p-value |
|---|---|---|---|---|
| CAP1 | 1 | 137.4 | 1.5333 | 0.02488* |
| Residual | 40 | 3584.1 | | |

**d**

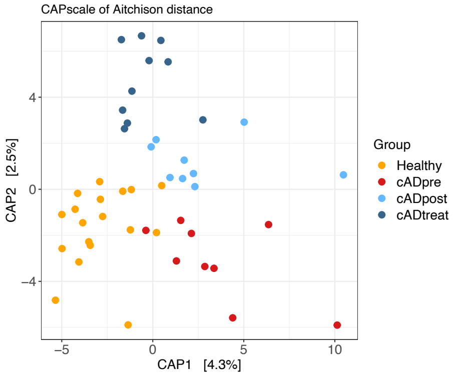


| | Df | Variance | F-statistic | p-value |
|---|---|---|---|---|
| Model | 3 | 291.4 | 1.1922 | 0.00557* |
| Residual | 42 | 3421.9 | | |

| | Df | Variance | F-statistic | p-value |
|---|---|---|---|---|
| Group | 3 | 291.4 | 1.1922 | 0.00584* |
| Residual | 42 | 3421.9 | | |

| | Df | Variance | F-statistic | p-value |
|---|---|---|---|---|
| CAP1 | 1 | 140 | 1.7188 | 0.00155* |
| Residual | 42 | 3421.9 | | |

**Fig. S4**

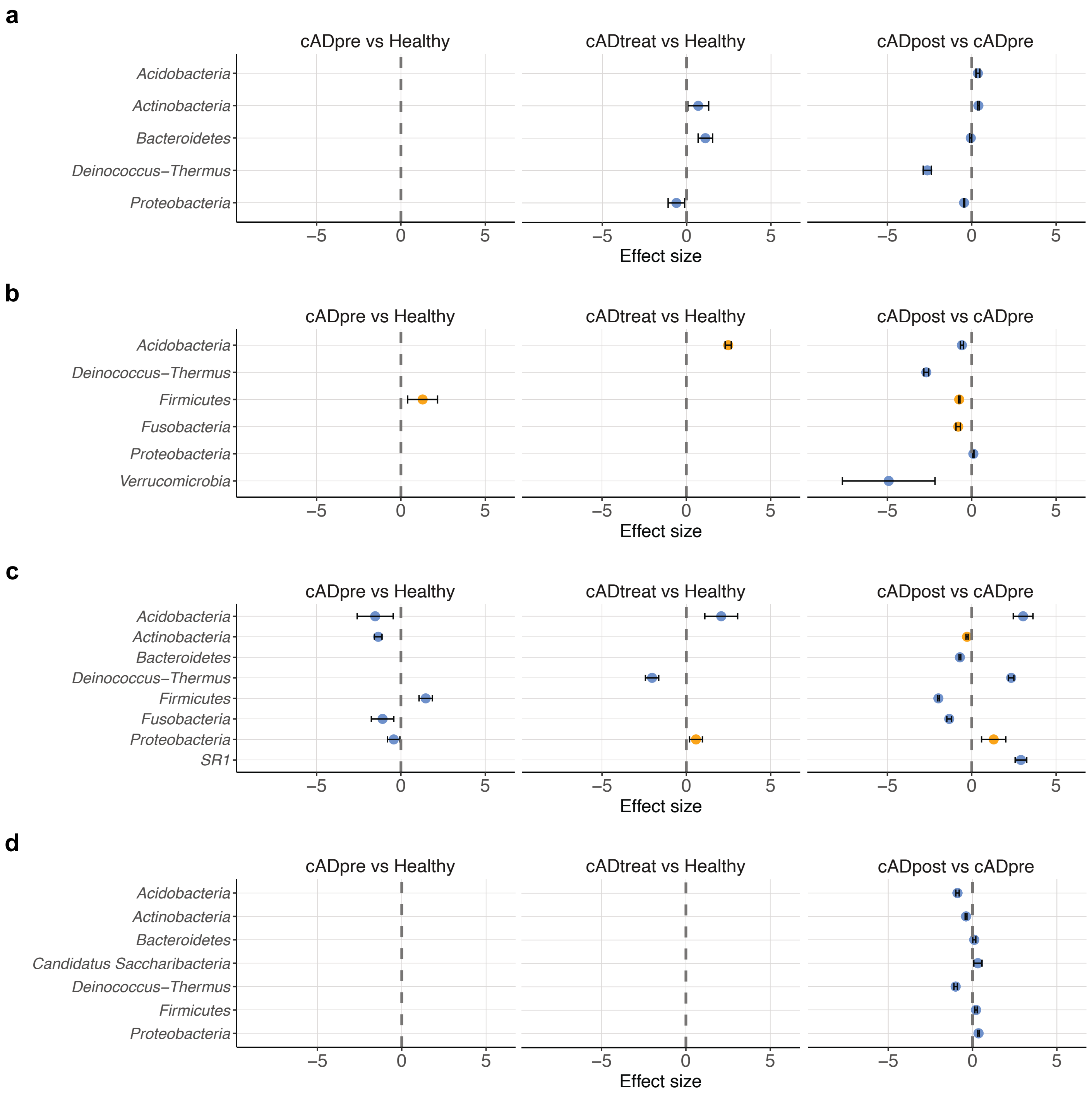
a
cADpre vs Healthy
cADtreat vs Healthy
cADpost vs cADpre
Acidobacteria
Actinobacteria
Bacteroidetes
Deinococcus–Thermus
Proteobacteria
–5
0
5
Effect size
b
cADpre vs Healthy
cADtreat vs Healthy
cADpost vs cADpre
Acidobacteria
Deinococcus–Thermus
Firmicutes
Fusobacteria
Proteobacteria
Verrucomicrobia
Effect size
c
cADpre vs Healthy
cADtreat vs Healthy
cADpost vs cADpre
Acidobacteria
Actinobacteria
Bacteroidetes
Deinococcus–Thermus
Firmicutes
Fusobacteria
Proteobacteria
SR1
Effect size
d
cADpre vs Healthy
cADtreat vs Healthy
cADpost vs cADpre
Acidobacteria
Actinobacteria
Bacteroidetes
Candidatus Saccharibacteria
Deinococcus–Thermus
Firmicutes
Proteobacteria
Effect size

**Fig. S5**

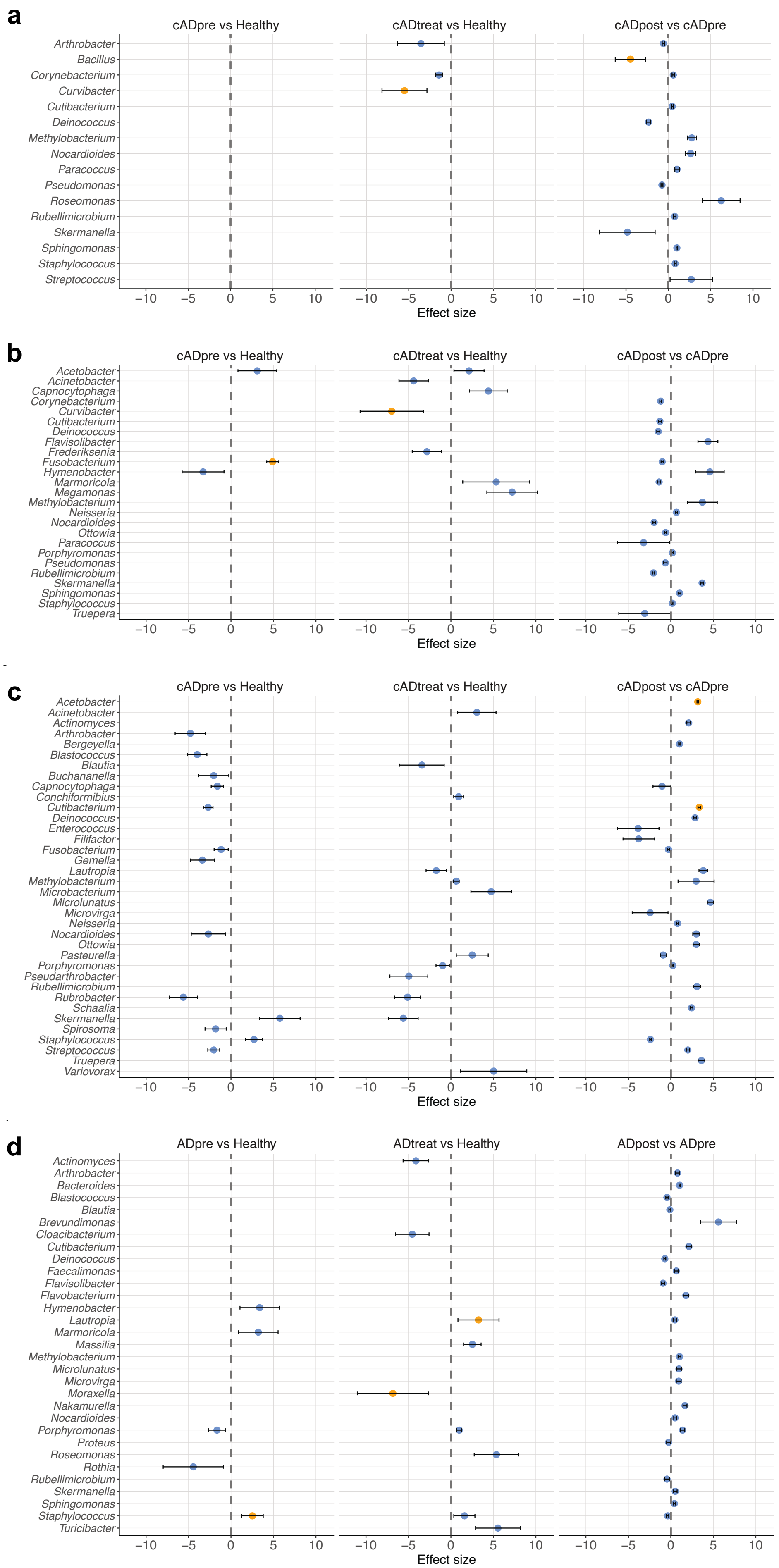
a
cADpre vs Healthy
cADtreat vs Healthy
cADpost vs cADpre
Arthrobacter
Bacillus
Corynebacterium
Curvibacter
Cutibacterium
Deinococcus
Methylobacterium
Nocardioides
Paracoccus
Pseudomonas
Roseomonas
Rubellimicrobium
Skermanella
Sphingomonas
Staphylococcus
Streptococcus
Effect size
b
cADpre vs Healthy
cADtreat vs Healthy
cADpost vs cADpre
Acetobacter
Acinetobacter
Capnocytophaga
Corynebacterium
Curvibacter
Cutibacterium
Deinococcus
Flavisolibacter
Frederiksenia
Fusobacterium
Hymenobacter
Marmoricola
Megamonas
Methylobacterium
Neisseria
Nocardioides
Ottowia
Paracoccus
Porphyromonas
Pseudomonas
Rubellimicrobium
Skermanella
Sphingomonas
Staphylococcus
Truepera
Effect size
c
cADpre vs Healthy
cADtreat vs Healthy
cADpost vs cADpre
Acetobacter
Acinetobacter
Actinomyces
Arthrobacter
Bergeyella
Blastococcus
Blautia
Buchananella
Capnocytophaga
Conchiformibius
Cutibacterium
Deinococcus
Enterococcus
Filifactor
Fusobacterium
Gemella
Lautropia
Methylobacterium
Microbacterium
Microlunatus
Microvirga
Neisseria
Nocardioides
Ottowia
Pasteurella
Porphyromonas
Pseudarthrobacter
Rubellimicrobium
Rubrobacter
Schaalia
Skermanella
Spirosoma
Staphylococcus
Streptococcus
Truepera
Variovorax
Effect size
d
ADpre vs Healthy
ADtreat vs Healthy
ADpost vs ADpre
Actinomyces
Arthrobacter
Bacteroides
Blastococcus
Blautia
Brevundimonas
Cloacibacterium
Cutibacterium
Deinococcus
Faecalimonas
Flavisolibacter
Flavobacterium
Hymenobacter
Lautropia
Marmoricola
Massilia
Methylobacterium
Microlunatus
Microvirga
Moraxella
Nakamurella
Nocardioides
Porphyromonas
Proteus
Roseomonas
Rothia
Rubellimicrobium
Skermanella
Sphingomonas
Staphylococcus
Turicibacter
Effect size

Fig. S6

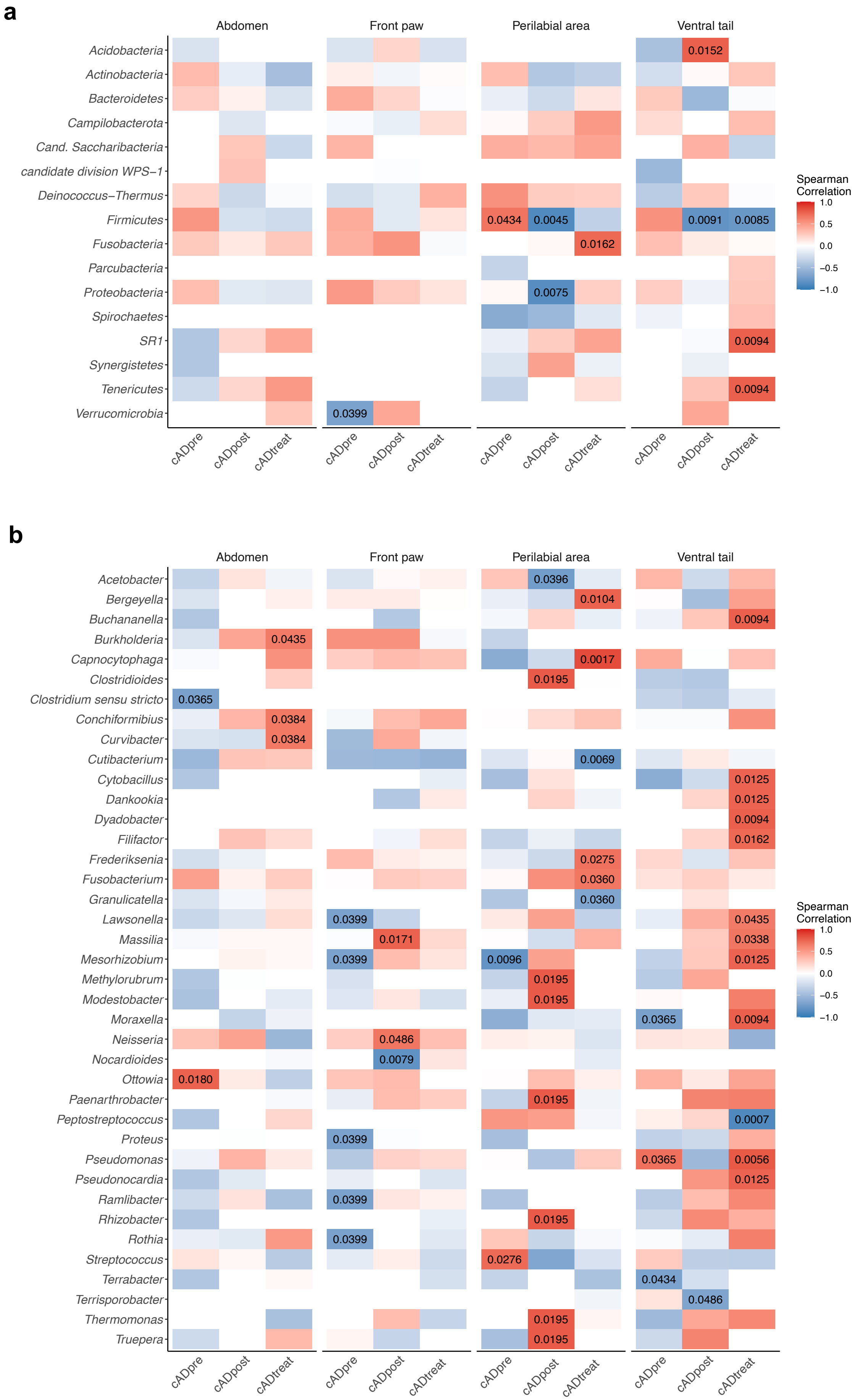
a
b
Abdomen
Front paw
Perilabial area
Ventral tail
Acidobacteria
Actinobacteria
Bacteroidetes
Campilobacterota
Cand. Saccharibacteria
candidate division WPS−1
Deinococcus−Thermus
Firmicutes
Fusobacteria
Parcubacteria
Proteobacteria
Spirochaetes
SR1
Synergistetes
Tenericutes
Verrucomicrobia
0.0152
0.0434
0.0045
0.0091
0.0085
0.0162
0.0075
0.0094
0.0094
0.0399
cADpre
cADpost
cADtreat
Spearman
Correlation
1.0
0.5
0.0
−0.5
−1.0
Acetobacter
Bergeyella
Buchananella
Burkholderia
Capnocytophaga
Clostridioides
Clostridium sensu stricto
Conchiformibius
Curvibacter
Cutibacterium
Cytobacillus
Dankookia
Dyadobacter
Filifactor
Frederiksenia
Fusobacterium
Granulicatella
Lawsonella
Massilia
Mesorhizobium
Methylorubrum
Modestobacter
Moraxella
Neisseria
Nocardioides
Ottowia
Paenarthrobacter
Peptostreptococcus
Proteus
Pseudomonas
Pseudonocardia
Ramlibacter
Rhizobacter
Rothia
Streptococcus
Terrabacter
Terrisporobacter
Thermomonas
Truepera

**Fig. S7**

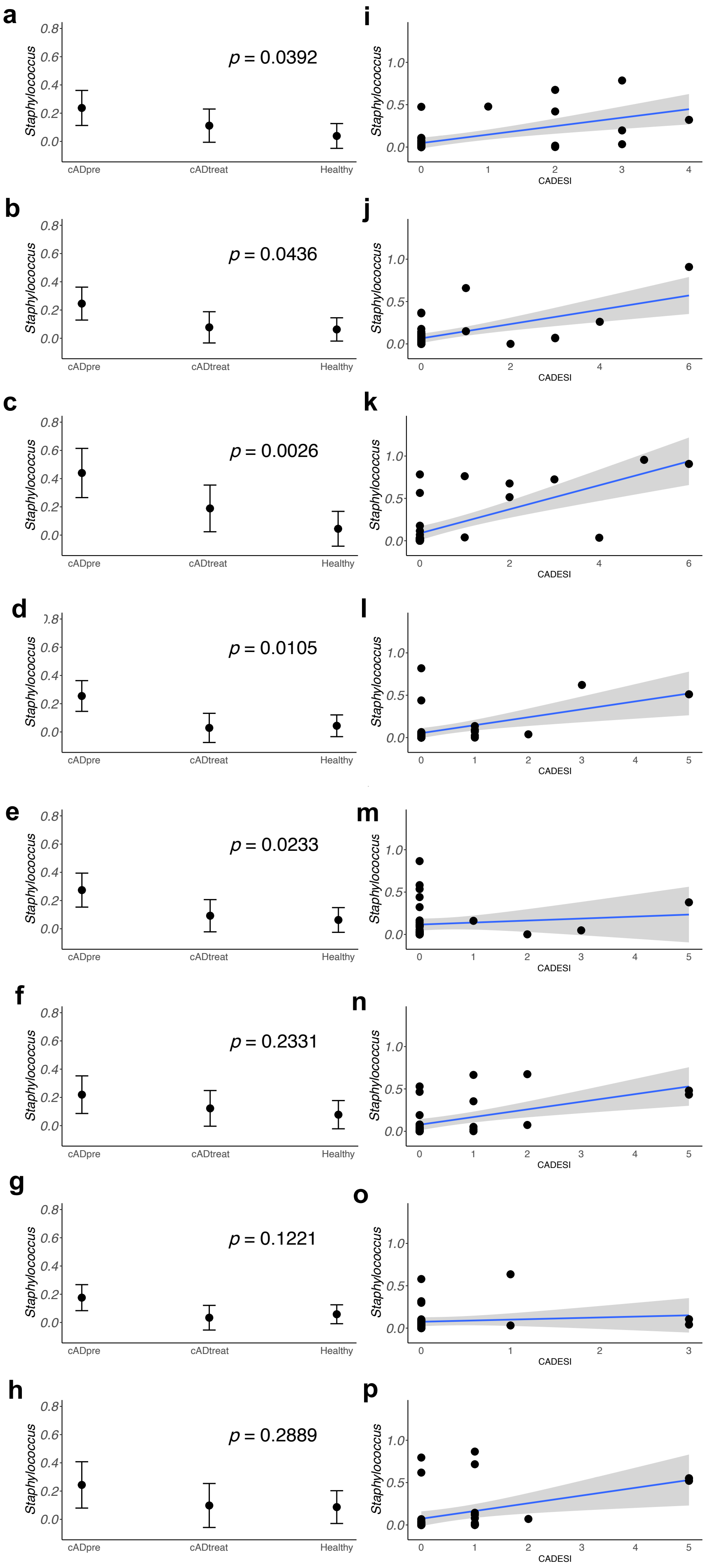
a
p = 0.0392
b
p = 0.0436
c
p = 0.0026
d
p = 0.0105
e
p = 0.0233
f
p = 0.2331
g
p = 0.1221
h
p = 0.2889
i
j
k
l
m
n
o
p
Staphylococcus
cADpre
cADtreat
Healthy
CADESI

**Fig. S8**

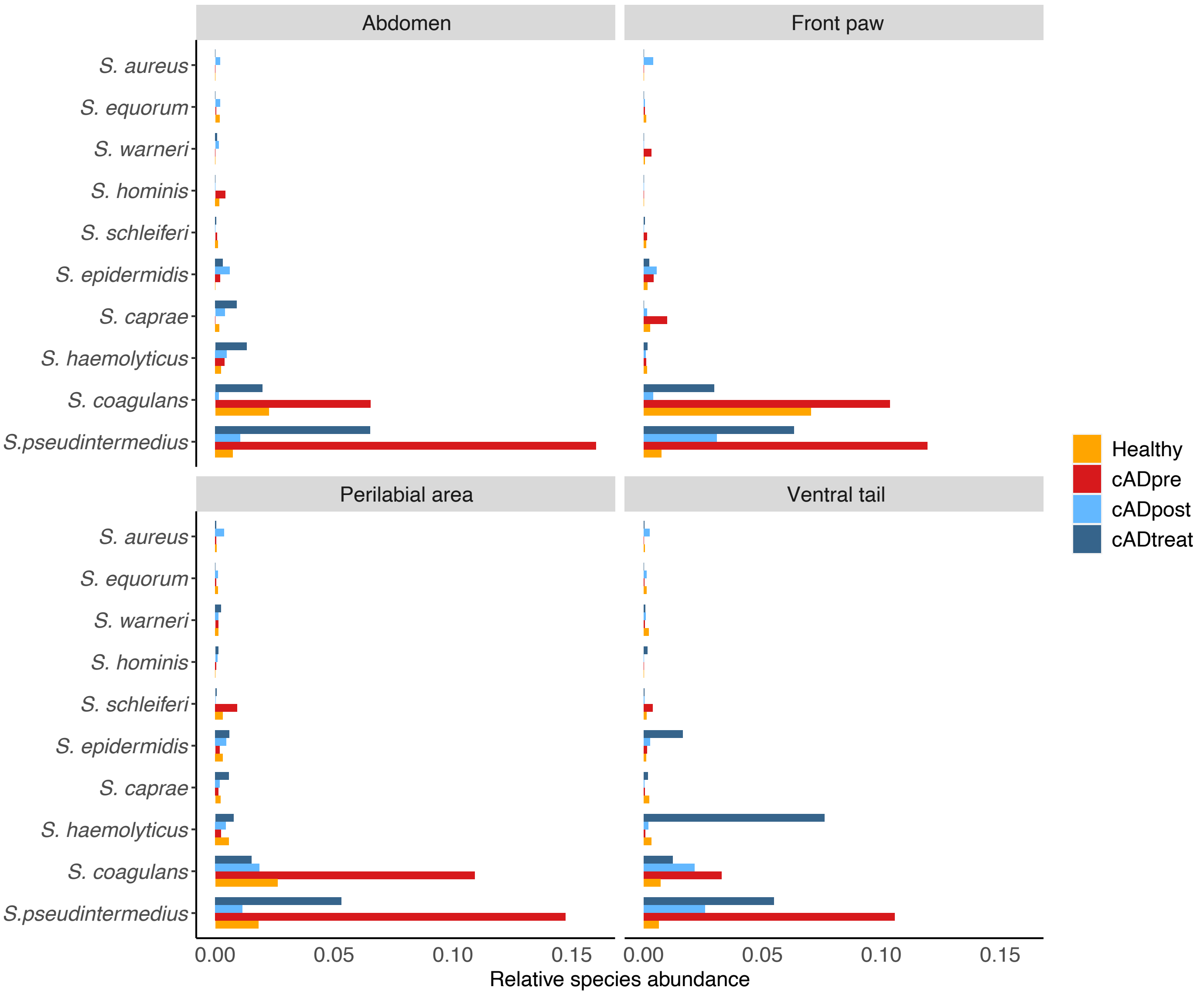

Abdomen
Front paw
Perilabial area
Ventral tail
S. aureus
S. equorum
S. warneri
S. hominis
S. schleiferi
S. epidermidis
S. caprae
S. haemolyticus
S. coagulans
S.pseudintermedius
0.00
0.05
0.10
0.15
Relative species abundance
Healthy
cADpre
cADpost
cADtreat

**Fig. S9**

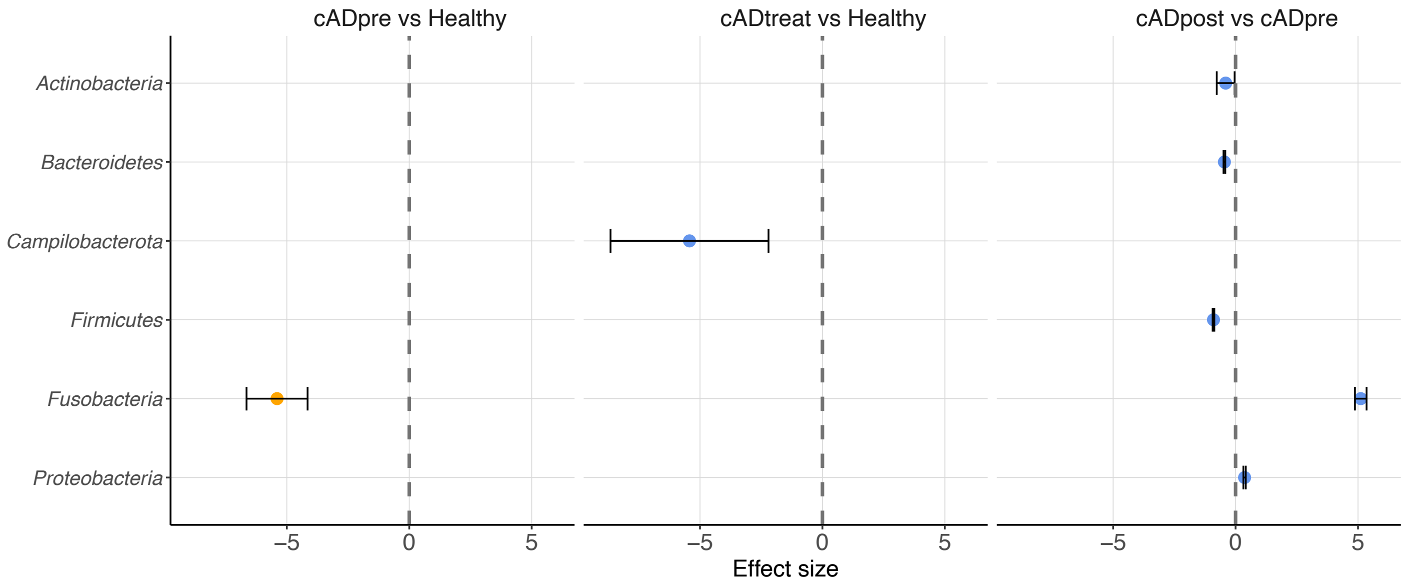

cADpre vs Healthy
cADtreat vs Healthy
cADpost vs cADpre
Actinobacteria
Bacteroidetes
Campilobacterota
Firmicutes
Fusobacteria
Proteobacteria
−5
0
5
Effect size

### Figure S10

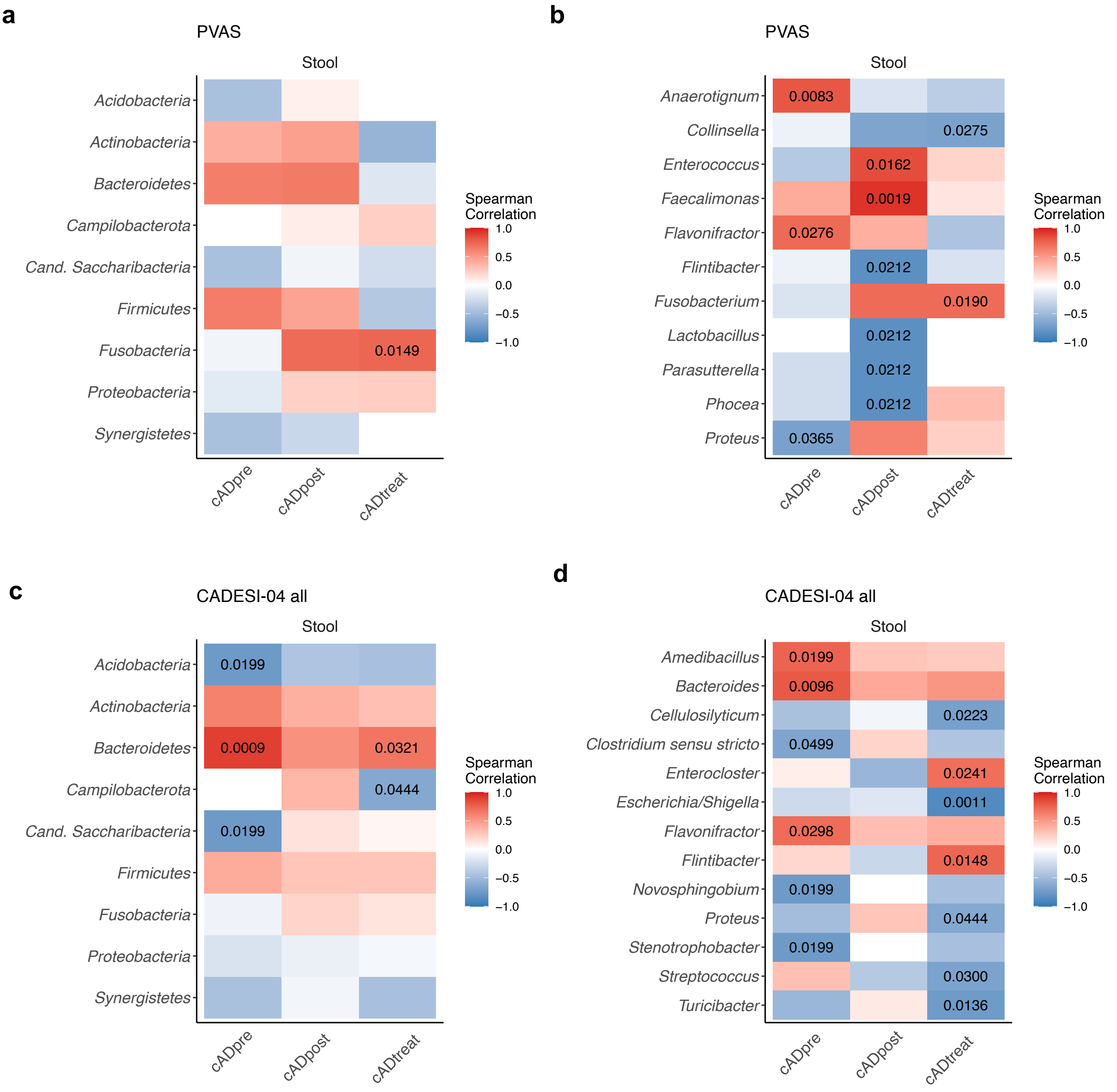
a
PVAS
Stool
Acidobacteria
Actinobacteria
Bacteroidetes
Campilobacterota
Cand. Saccharibacteria
Firmicutes
Fusobacteria
Proteobacteria
Synergistetes
0.0149
cADpre
cADpost
cADtreat
Spearman Correlation
1.0
0.5
0.0
−0.5
−1.0
b
PVAS
Stool
Anaerotignum
Collinsella
Enterococcus
Faecalimonas
Flavonifractor
Flintibacter
Fusobacterium
Lactobacillus
Parasutterella
Phocea
Proteus
0.0083
0.0275
0.0162
0.0019
0.0276
0.0212
0.0190
0.0212
0.0212
0.0212
0.0365
cADpre
cADpost
cADtreat
Spearman Correlation
1.0
0.5
0.0
−0.5
−1.0
c
CADESI-04 all
Stool
Acidobacteria
Actinobacteria
Bacteroidetes
Campilobacterota
Cand. Saccharibacteria
Firmicutes
Fusobacteria
Proteobacteria
Synergistetes
0.0199
0.0009
0.0321
0.0444
0.0199
cADpre
cADpost
cADtreat
Spearman Correlation
1.0
0.5
0.0
−0.5
−1.0
d
CADESI-04 all
Stool
Amedibacillus
Bacteroides
Cellulosilyticum
Clostridium sensu stricto
Enterocloster
Escherichia/Shigella
Flavonifractor
Flintibacter
Novosphingobium
Proteus
Stenotrophobacter
Streptococcus
Turicibacter
0.0199
0.0096
0.0223
0.0499
0.0241
0.0011
0.0298
0.0148
0.0199
0.0444
0.0199
0.0300
0.0136
cADpre
cADpost
cADtreat
Spearman Correlation
1.0
0.5
0.0
−0.5
−1.0

### Figure S11

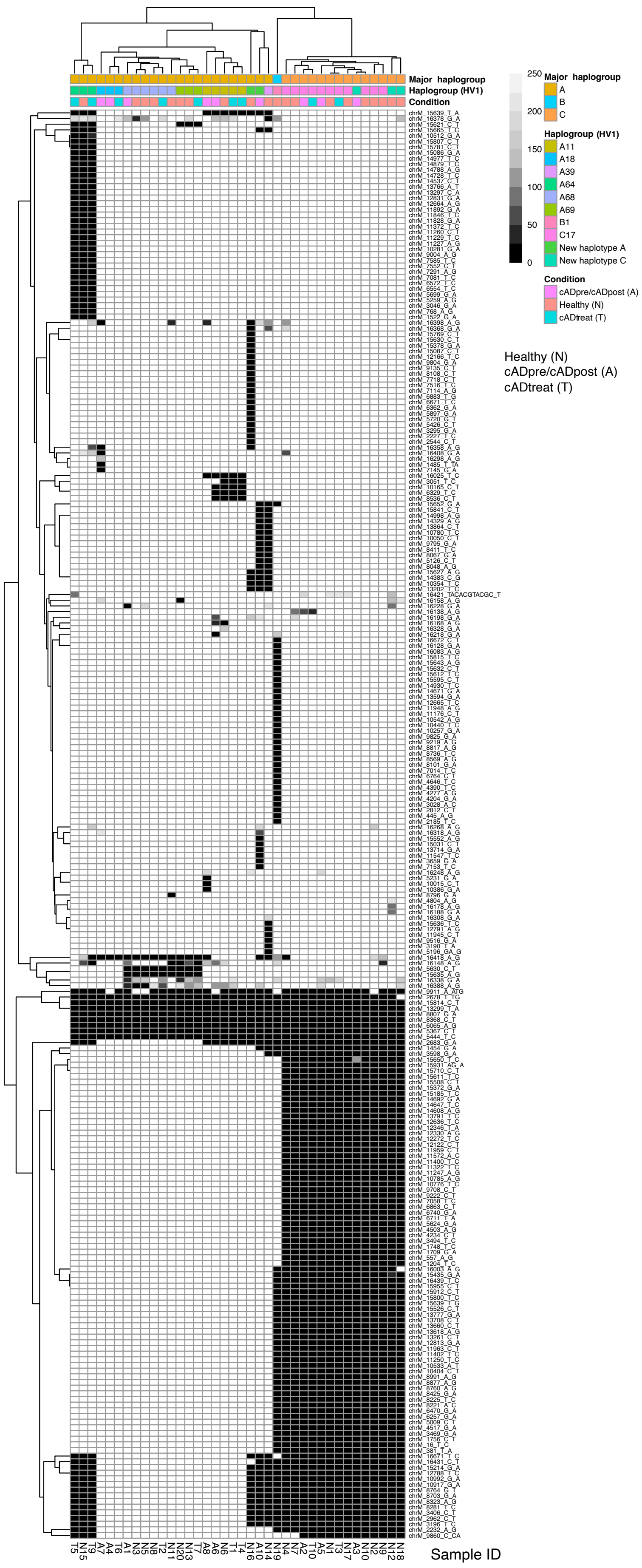

Major haplogroup
Haplogroup (HV1)
Condition
250
200
150
100
50
0
Major haplogroup
A
B
C
Haplogroup (HV1)
A11
A18
A39
A64
A68
A69
B1
C17
New haplotype A
New haplotype C
Condition
cADpre/cADpost (A)
Healthy (N)
cADtreat (T)
Healthy (N)
cADpre/cADpost (A)
cADtreat (T)
Sample ID
